# A unique view of SARS-CoV-2 through the lens of ORF8 protein

**DOI:** 10.1101/2020.08.25.267328

**Authors:** Sk. Sarif Hassan, Shinjini Ghosh, Diksha Attrish, Pabitra Pal Choudhury, Murat Seyran, Damiano Pizzol, Parise Adadi, Tarek Mohamed Abd El-Aziz, Antonio Soares, Ramesh Kandimalla, Kenneth Lundstrom, Murtaza Tambuwala, Alaa A. A. Aljabali, Amos Lal, Gajendra Kumar Azad, Vladimir N. Uversky, Samendra P. Sherchan, Wagner Baetas-da-Cruz, Bruce D. Uhal, Nima Rezaei, Adam M. Brufsky

**Affiliations:** Department of Mathematics, Pingla Thana Mahavidyalaya, Maligram 721140, India; Department of Biophysics, Molecular Biology and Bioinformatics, University of Calcutta, Kolkata 700009, West Bengal, India; Dr. B. R. Ambedkar Centre For Biomedical Research (ACBR), University of Delhi (North Campus), Delhi 110007, India; Applied Statistics Unit, Indian Statistical Institute, Kolkata 700108, West Bengal, India; Doctoral studies in natural and technical sciences (SPL 44), University of Vienna, Austria; Italian Agency for Development Cooperation - Khartoum, Sudan Street 33, Al Amarat, Sudan; Department of Food Science, University of Otago, Dunedin 9054, New Zealand; Department of Cellular and Integrative Physiology, University of Texas Health Science Center at San Antonio, 7703 Floyd Curl Dr, San Antonio, TX 78229-3900, USA & Zoology Department, Faculty of Science, Minia University, El-Minia 61519, Egypt; CSIR-Indian Institute of Chemical Technology Uppal Road, Tarnaka, Hyderabad-500007, Telangana State, India; PanTherapeutics, Rte de Lavaux 49, CH1095 Lutry, Switzerland; School of Pharmacy and Pharmaceutical Science, Ulster University, Coleraine BT52 1SA, Northern Ireland, UK; Department of Pharmaceutics and Pharmaceutical Technology, Yarmouk University-Faculty of Pharmacy, Irbid 566, Jordan; Division of Pulmonary and Critical Care Medicine, Mayo Clinic, Rochester, Minnesota, USA; Department of Zoology, Patna University, Patna-800005, Bihar, India; Department of Molecular Medicine, Morsani College of Medicine, University of South Florida, Tampa, FL 33612, USA; Department of Environmental Health Sciences, Tulane University, New Orleans, LA, 70112, USA; Translational Laboratory in Molecular Physiology, Centre for Experimental Surgery, College of Medicine, Federal University of Rio de Janeiro (UFRJ), Rio de Janeiro, Brazil; Department of Physiology, Michigan State University, East Lansing, MI 48824, USA; Research Center for Immunodeficiencies, Pediatrics Center of Excellence, Children’s Medical Center, Tehran University of Medical Sciences, Tehran, Iran & Network of Immunity in Infection, Malignancy and Autoimmunity (NIIMA), Universal Scientific Education and Research Network (USERN), Stockholm, Sweden; University of Pittsburgh School of Medicine, Department of Medicine, Division of Hematology/Oncology, UPMC Hillman Cancer Center, Pittsburgh, PA, USA

**Keywords:** ORF8, SARS-CoV-2, Mutations, COVID-19 and Genetic variations

## Abstract

Immune evasion is one of the unique characteristics of COVID-19 attributed to the ORF8 protein of severe acute respiratory syndrome coronavirus 2 (SARS-CoV-2). This protein is involved in modulating the host adaptive immunity through downregulating MHC (Major Histocompatibility Complex) molecules and innate immune responses by surpassing the interferon mediated antiviral response of the host. To understand the immune perspective of the host with respect to the ORF8 protein, a comprehensive study of the ORF8 protein as well as mutations possessed by it, is performed. Chemical and structural properties of ORF8 proteins from different hosts, that is human, bat and pangolin, suggests that the ORF8 of SARS-CoV-2 and Bat RaTG13-CoV are very much closer related than that of Pangolin-CoV. Eighty-seven mutations across unique variants of ORF8 (SARS-CoV-2) are grouped into four classes based on their predicted effects. Based on geolocations and timescale of collection, a possible flow of mutations was built. Furthermore, conclusive flows of amalgamation of mutations were endorsed upon sequence similarity and amino acid conservation phylogenies. Therefore, this study seeks to highlight the uniqueness of rapid evolving SARS-CoV-2 through the ORF8.

## 1. Introduction

Severe acute respiratory syndrome-coronavirus-2 (SARS-CoV-2) is a novel coronavirus whose first outbreak was reported in December 2019 in Wuhan, China, where a cluster of pneumonia cases was detected, and on 11th March, 2020, WHO declared this outbreak a pandemic [1, 2, 3]. As of 22nd August 2020, a total of 22.9 million confirmed COVID-19 cases have been reported worldwide with 7,99,350 deaths [4]. SARS-CoV-2 belongs to the family Coronaviridae and has 55% nucleotide similarity and 30% protein sequence similarity with SARS-CoV, which caused the previous outbreak of SARS in 2002 [5, 6]. SASR-CoV2 is a single-stranded RNA virus of positive polarity whose genome is approximately 30 kb in length and encodes for 16 non-structural proteins, four structural and six accessory proteins [7, 8]. The SARS-CoV-2 has a total of six accessory proteins including 3a, 6, 7a, 7b, 8 and 10 [9, 10]. Among these accessory proteins, ORF8 is a significantly exclusive protein as it is different from another known SARS-CoV and thus associated with high efficiency in pathogenicity transmission [11, 12]. The SARS-CoV-2 ORF8 displays arrays of functions; inhibition of interferon 1, promoting viral replication, inducing apoptosis and modulating ER stress [13, 14, 15].

The SARS-CoV-2 ORF8 is a 121 amino acid long protein, which has an N-terminal hydrophobic signal peptide (1-15 aa) and an ORF8 chain (16-121 aa) [16, 17]. The functional motif (VLVVL) of SARS-CoV-ORF8b, which is responsible for induction of cell stress pathways and activation of macrophages is absent from the SARS-CoV-2 ORF8 protein [18]. In the later stages of the SARS-CoV epidemic it was found that a 29 nucleotide deletion in the ORF8 protein caused it to split into ORF8a (39 aa) and ORF8b (84aa) rendering it functionless while the SARS-CoV-2 ORF8 is intact [19]. Also, the SARS-CoV ORF8 had a function in interspecies transmission and viral replication efficiency as a reported 382 nucleotide deletion, which included ORF8ab resulted in a reduced ability of viral replication in human cells [20]. However, the SARS-CoV-2 ORF8 mainly acts as an immune-modulator by down-regulating MHC class I molecules, therefore protecting the infected cells against cytotoxic T cell killing of target cells. Simultaneously it was proposed forth that it is a potential inhibitor of type 1 interferon signalling pathway which is a key component of antiviral host immune response [21, 22]. The ORF8 also regulates unfolded protein response (UPR) induced due to ER stress by triggering ATF-6 activation, thus promoting infected cell survivability for its own benefit [23]. Since this protein impacts various host pathogen processes and developed various strategies that allow it to escape through host immune responses, it becomes important to study the mutations in the ORF8 to develop a better understanding of the viral infectivity and for developing efficient therapeutic drugs against it [24].

In this present study, we identified the distinct mutations present across unique variants of the SARS-CoV-2 ORF8 and classified them according to their predicted effect on the host, i.e disease or neutral and the consequences on protein structural stability. Furthermore, we compared the ORF8 protein of SARS-CoV-2 with that of Bat-RaTG13-CoV and Pangolin-CoV ORF8 and tried to determine the evolutionary relationships with respect to sequence similarity and discussed the rising concern on its originality. In addition we made a study on polarity and charge of the SARS-CoV-2 ORF8 mutations in two of its distinct domains and explored the possible effect on functionality changes. Following this, we present the possible flow of mutations considering different geographical locations and chronological time scale simultaneously, validating with sequence-based and amino acid conservation-based phylogeny and thereby predicted the possible route taken through the course of assimilation of mutations.

## 2. Materials and Methods

### 2.1. Data: ORF8 of SARS-CoV-2

As of 14th August, 2020, there were 10,314 complete genomes of SARS-CoV-2 available in the NCBI database and accordingly each genome contains one of the accessory proteins ORF8 and among them only 127 sequences were found to be unique. The amino acid sequences of the ORF8 were exported in fasta format using file operations through Matlab [25]. Among these 127 unique ORF8 protein sequences, only 96 sequences possess various mutations and the remaining sequences do not either possess any mutations or possess ambiguous mutations. In this present study, we only concentrate on 96 ORF8 proteins, which are listed in Table 1. In order to find mutations, we hereby consider the reference ORF8 protein as the ORF8 sequence (YP_009724396.1) of the SARS-CoV-2 genome (NC_045512) from China: Wuhan [26].

**Table 1:**
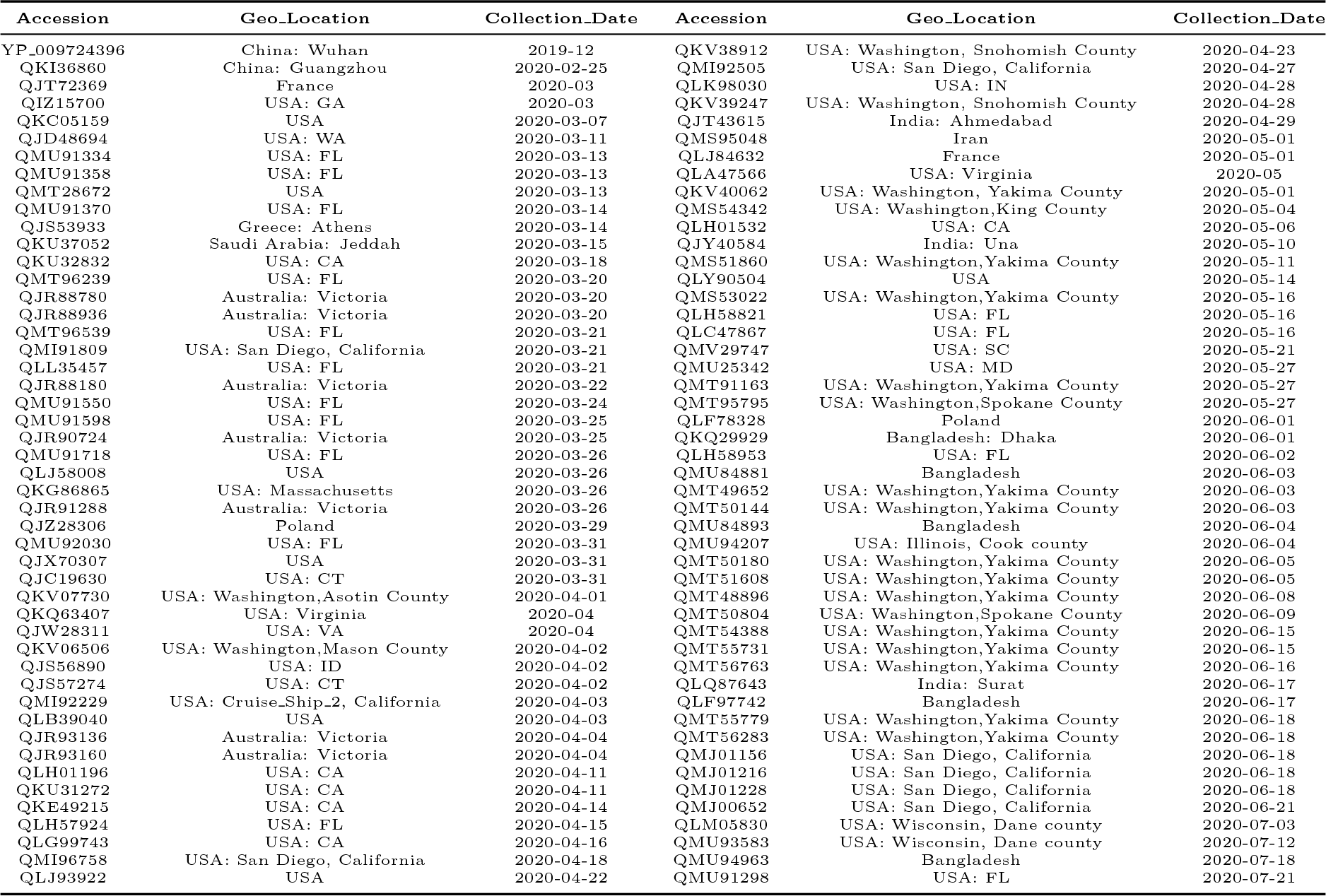
96 unique ORF8 protein IDs with associated information

In addition, the ORF8 protein sequences of SARS-CoV, Pangolin and Bat RaTG13-CoVs were retrieved from the NCBI as reference sequences for understanding the evolutionary proximal origin of the protein ORF8 of SARS-CoV-2. Note that, only one unique ORF8 variant is obtained among four available sequences (QIA48620.1, QIA48638.1, QIA48647.1, QIQ54055.1) of Pangolin-CoV and therefore the ORF8 protein of Pangolin-CoV is absolutely conserved. There are three available ORF8 (AVP78048.1, AVP78037.1, QHR63307.1 (Bat RatG13)) of Bat-CoV and among them two variants (QHR63307.1 (Bat RatG13)) and AVP78048.1) are turned out to be different with 96% sequence similarity. The alignment is shown in Fig. 1.

**Figure 1:**
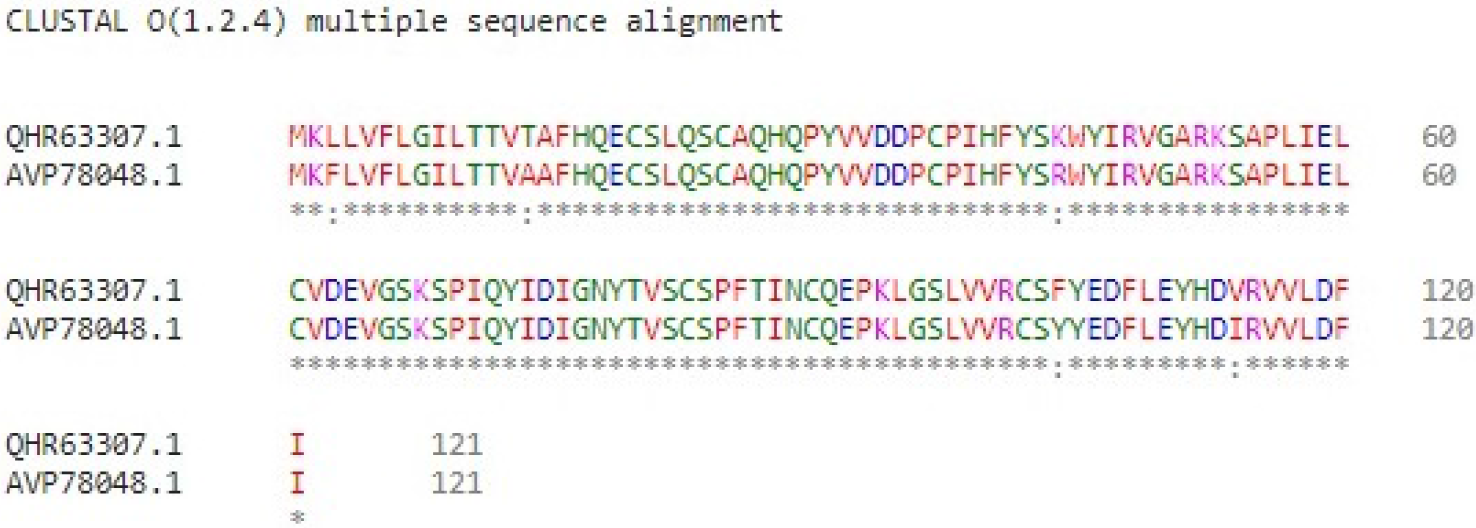
Alignment of two ORF8 sequences (116 is identical residues among 121) of Bat-CoV

Only the mutations L3F, T14A, K44R, F104Y and V114I are embedded in the ORF8 proteins of Bat-CoV.

### 2.2. Methods

#### 2.2.1. Mutation Identification

Mutations in proteins are responsible for several genetic orders/disorders. Identifying these mutations requires novel detection methods, which have been reported in the literature [27]. In this study, each unique ORF8 sequence was aligned using NCBI protein p-blast and sometimes using omega blast suites to determine the mismatches and thereby the missense mutations (amino acid changes) are identified [28, 29]. For the effect of identified mutations, a web-server ‘Meta-SNP’ was used and also for the structural effects of mutations, another web-server ‘I-MUTANT’ was used [30, 31]. The web-server ‘QUARK’ was used for prediction of the secondary structure of ORF8 proteins [32, 33].

#### 2.2.2. Amino Acids Compositions and Phylogeny

A protein sequence of ORF8 is composed of twenty different amino acids with various frequencies. The frequency of occurrence of each amino acid *A_i_* is determined for each primary sequence of ORF8 proteins. Hence for each of the 96 ORF8 proteins, a twenty dimensional vector considering the frequency of twenty amino acids can be obtained. A distance matrix (Euclidean distance) is formed by determining the distances (pairwise) among the twenty dimensional frequency vectors for each ORF8 protein [34, 35]. Thereby, applying the nearest neighbour joining method, a phylogeny is derived using the distance matrix formed by the twenty dimensional frequency vectors for each of the ORF8 proteins of interest [36, 37, 38, 39].

#### 2.2.3. Evaluating the propensity of various ORF8 proteins for intrinsic disorder

Per-residue disorder distribution in sequences of query proteins was evaluated by PONDR-VSL2 [40], which is one of the more accurate standalone disorder predictors [41, 42, 43]. The per-residue disorder predisposition scores are on a scale from 0 to 1, where values of 0 indicate fully ordered residues, and values of 1 indicate fully disordered residues. Values above the threshold of 0.5 are considered disordered residues, whereas residues with disorder scores between 0.25 and 0.5 are considered highly flexible, and residues with disorder scores between 0.1 and 0.25 are taken as moderately flexible.

## 3. Results

### 3.1. ORF8 of SARS-CoV-2

The SARS-CoV-2 ORF8 protein (YP_009724396) is a 121 amino acid long protein which has an N-terminal hydrophobic signal peptide (1-15 aa) and an ORF8 chain (16-121 aa). A schematic representation of ORF8 (SARS-CoV-2) is presented in Fig. 2.

**Figure 2:**
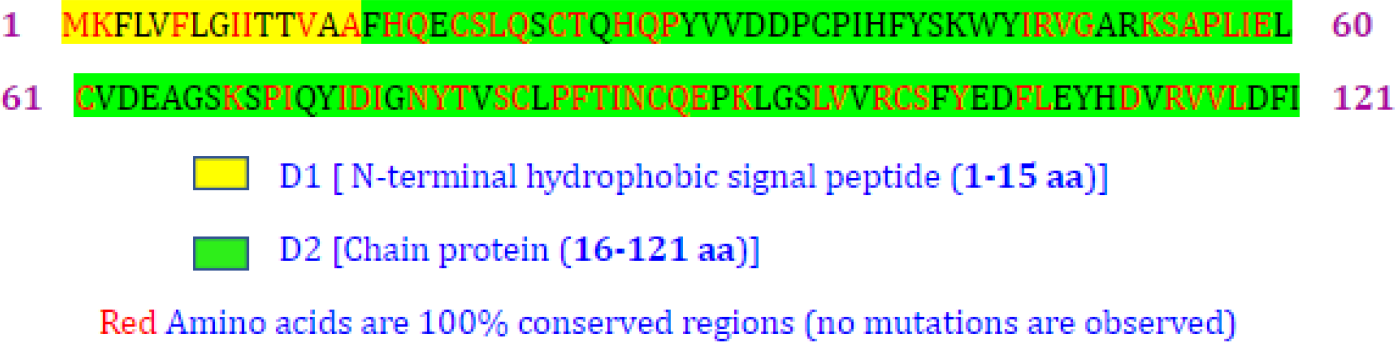
A schematic representation of sub-domains and conserved amino acid residues of the ORF8 protein of SARS-CoV-2

It was observed that the total number (63) of hydrophilic residues is more than that (58) of the hydrophobic residues. However, from the predicted secondary structure (Fig. 3), it was found that the highest solubility score is four, indicating that although hydrophilic residues are higher in number, however, they are not highly exposed to the external environment since they are folded inside of the protein therefore this protein is poorly soluble.

**Figure 3:**
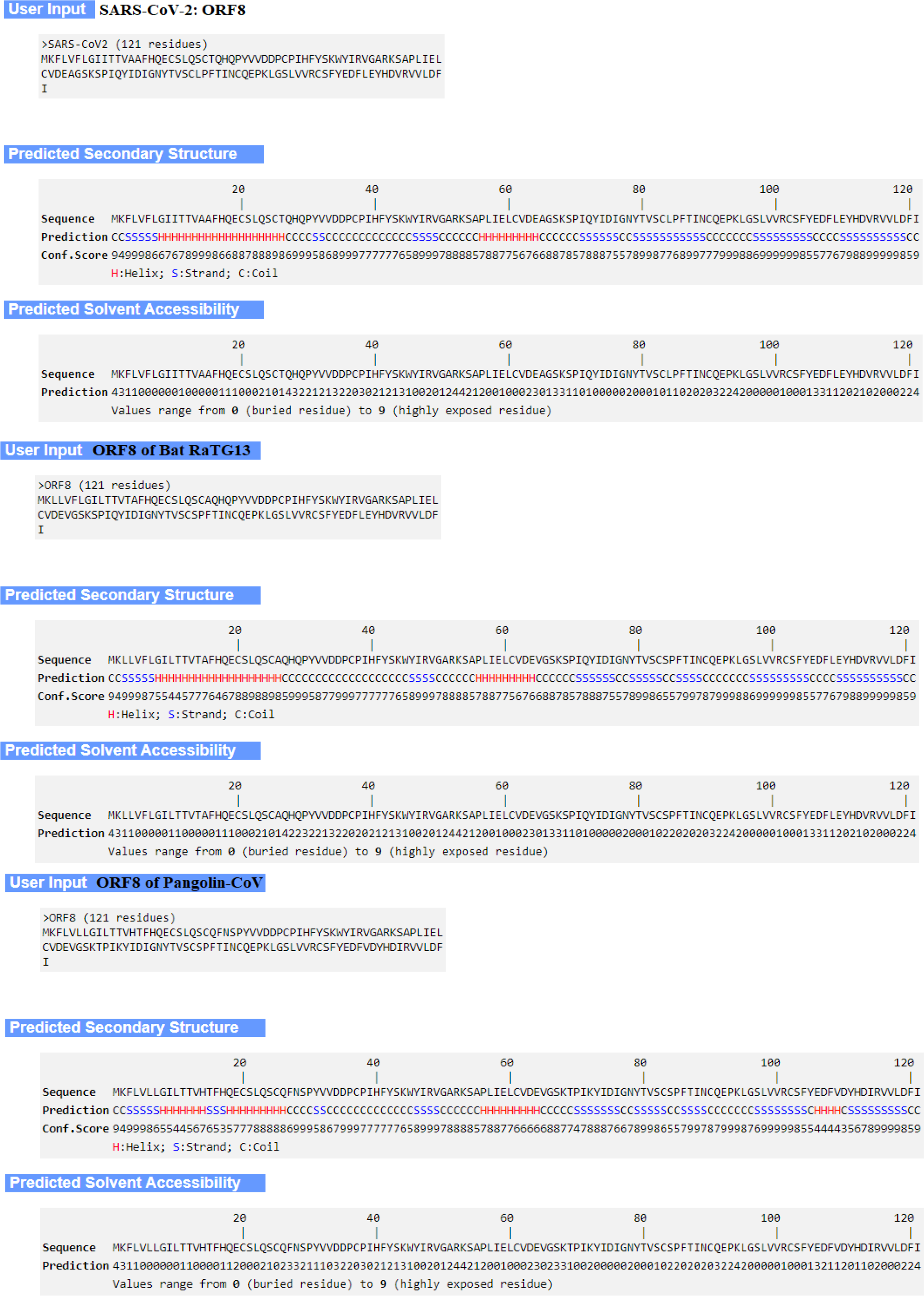
Predicted secondary and solvent accessibility of the ORF8 of SARS-CoV-2(top), Bat RaTG13-CoV(middle) and Pangolin-CoV (bottom)

We further predicted the secondary structure as well as solvent accessibility of ORF8 proteins of SARS-CoV-2, Bat-RaTG13-CoV and Pangolin-CoV using the ab-initio webserver QUARK (Fig. 3) and tried to perceive the differences.

Comparing the ORF8 secondary structures of SARS-CoV-2, Bat-RaTG13-CoV and Pangolin-CoV, the following changes are found at four different locations as presented in Table 2.

**Table 2:** Changes of secondary sub-structures in ORF8 among the three hosts

| aa positions | ORF8 (SARS-CoV-2) | ORF8 (Bat RatG13-CoV) | ORF8 (Pangolin-CoV) |
| --- | --- | --- | --- |
| 15-17 | HHH | HHH | SSS |
| 31-32 | SS | CC | SS |
| 84-85 | SS | CC | CC |
| 106-109 | CCCC | CCCC | HHHH |

From Table 2, it is inferred that the secondary structures of ORF8 (SARS-CoV-2) and that of Bat-RaTG13 are much closer related than the ORF8 of Pangolin-CoV.

It was obtained that the largest conserved region in the ORF8 protein of SARS-CoV-2 is ‘PFTINCQE’ (in D2), eight amino acids long (Table 3) as seen in Fig. 4. It turned out that the 6.6% region is 100% evolutionary conserved across all 96 distinct variants of the 121 amino acid long ORF8 protein.

**Figure 4:**
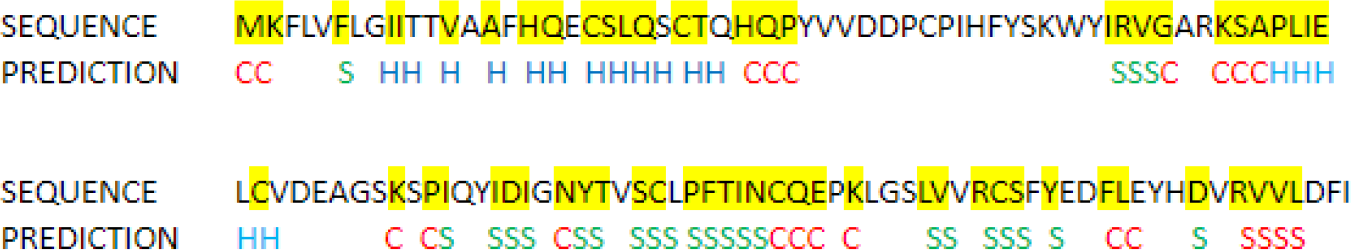
Conserved regions (marked in yellow) and predicted secondary structure of the ORF8 protein of SARS-CoV-2

**Table 3:** Conserved regions in the ORF8 of SARS-CoV-2

| Conserved region | Length | Domain | Conserved region | Length | Domain | Conserved region | Length | Domain |
| --- | --- | --- | --- | --- | --- | --- | --- | --- |
| MK | 2 | D1 | HQP | 3 | D2 | SC | 2 | D2 |
| F | 1 | D1 | IRVG | 4 | D2 | **PFTINCQE** | **8** | **D2** |
| II | 2 | D1 | KSAPLIE | 7 | D2 | K | 1 | D2 |
| V | 1 | D1 | C | 1 | D2 | LV | 2 | D2 |
| A | 1 | D1 | K | 1 | D2 | RCS | 3 | D2 |
| HQ | 2 | D2 | PI | 2 | D2 | Y | 1 | D2 |
| CSLQ | 4 | D2 | IDI | 3 | D2 | FL | 2 | D2 |
| CT | 2 | D2 | NYT | 3 | D2 | D | 1 | D2 |
|  |  |  |  |  |  | RVVL | 4 | D2 |

Most of the conserved regions in SARS-CoV-2 ORF8 (Table 3 and Fig. 4) lie around helix-coil and strand-coil junctions signifying the functional importance of these regions. Helix regions also have conserved amino acids. It can be hypothesised that these junctions are involved in protein-protein interactions. Therefore, these regions are conserved in nature.

The ORF8 protein of SARS-CoV-2 has only 55.4% nucleotide similarity and 30% protein identity with that of SARS-CoV as shown in Fig. 5.

**Figure 5:**
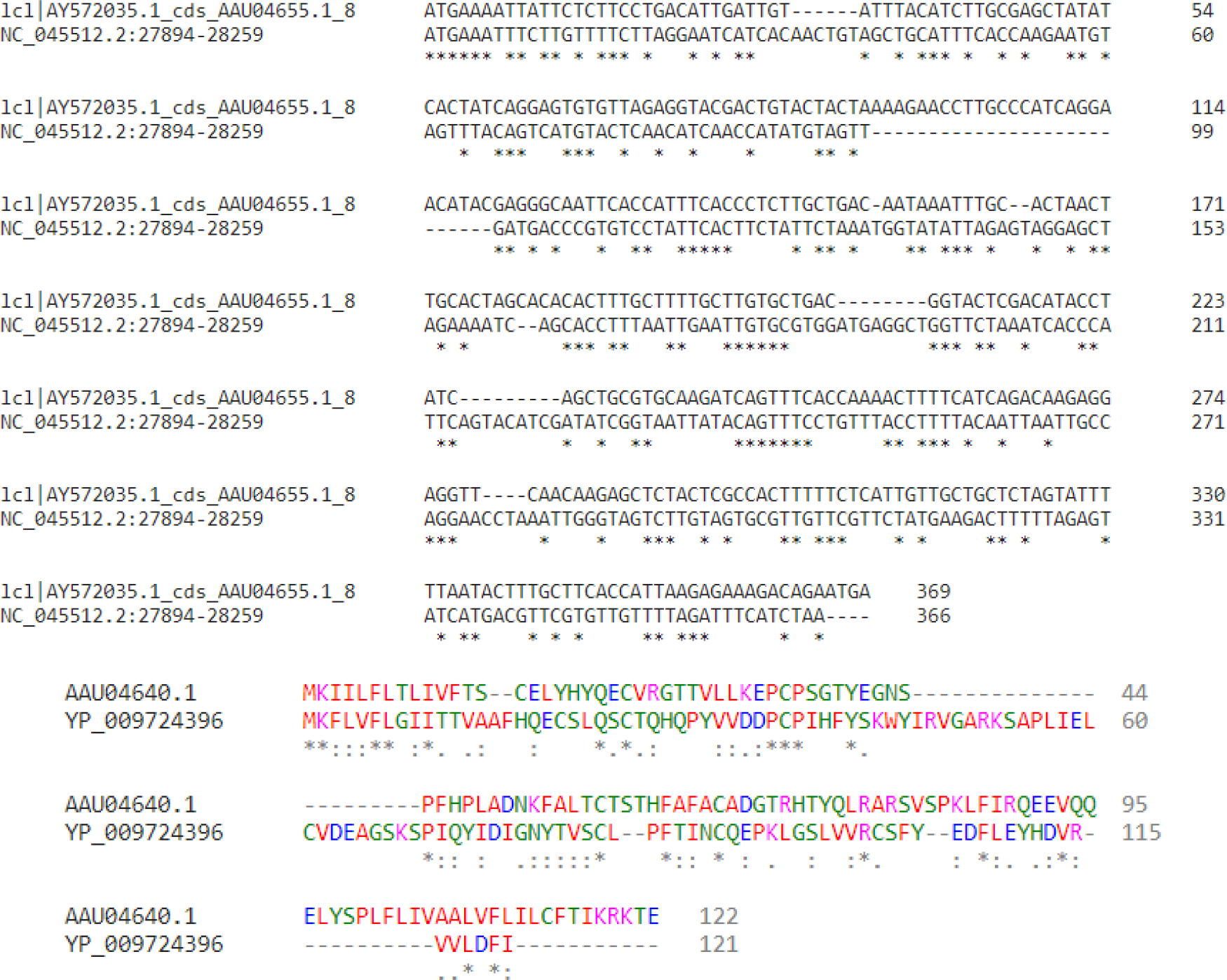
Sequence alignment of ORF8 protein of SARS-CoV and SARS-CoV-2

Although the SARS-CoV-2 ORF8 has differentiating genome characteristics but exhibit high functional similarity with that of SARS-CoV ORF8ab:

- The SARS-CoV ORF8ab original protein was found to have an N-terminal hydrophobic signal sequence that directs its transport to the endoplasmic reticulum (ER). However, after deletion of 29 nucleotides, which split the ORF8ab protein into ORF8a and ORF8b, it was found that only ORF8a was able to translocate to the ER while ORF8b remained distributed throughout the cell. Likewise, the SARS-CoV-2 ORF8 protein also contains an N-terminal hydrophobic signal peptide (1-15 aa), which is involved in the same function.
- The ER has an internal oxidative environment akin to other organelles, which is necessary for protein folding and oxidation processes. Due to this oxidative environment, formation of intra or inter-molecular cysteine bonds between unpaired cysteine residues takes place as the SARS-CoV ORF8ab protein is an ER resident protein and there are ten cysteine residues present, which exhibit disulphide linkages and form homomultimeric complexes within the ER. Similarly, the ORF8 of SARS-CoV-2 has also seven cysteines, which may be expected to form these types of disulphide linkages.
- The SARS-CoV ORF8ab is characterized by the presence of an asparagine residue at position 81 with the Asn-Val-Thr motif responsible for N-linked glycosylation, whereas SARS-CoV-2 ORF8 has an N-linked glycosylation site at Asn (78) and the motif is Asn-Tyr-Thr.
- The SARS-CoV-2 ORF8 is found to have both protein-protein and protein-DNA interactions while SARS-CoV ORF8ab shows only protein-protein interactions.

Further, it was found that the ORF8 protein of SARS-CoV-2 is very much similar (95%) to that of Bat-CoV RaTG13 on the basis of sequence similarity as well as on the basis of phylogenetic relationships as shown in Fig. 6.

**Figure 6:**
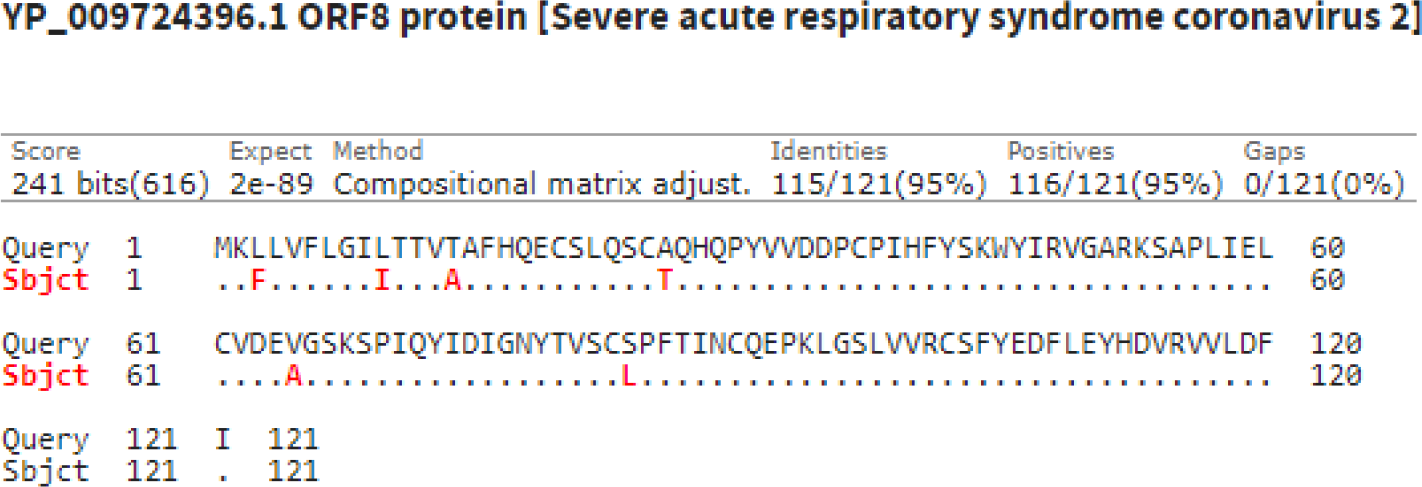
Amino acid sequence alignment of ORF8 proteins of Bat RaTG13 and SARS-CoV-2

As we can see from the sequence alignment, there are only six amino acid differences between the SARS-CoV-2 ORF8 and Bat-CoV RaTG13. All of these mutations in the ORF8 protein with respect to the reference ORF8 sequence of Bat-RaTG13 were found to be neutral type as predicted through webserver Meta-SNP and all of them had a decreasing effect on stability of the protein as determined using the server (I-MUTANT).

We have also aligned the Pangolin-CoV ORF8 sequence with that of SARS-CoV-2 and found that there is a sequence similarity of 88%, as depicted in Fig. 7.

**Figure 7:**
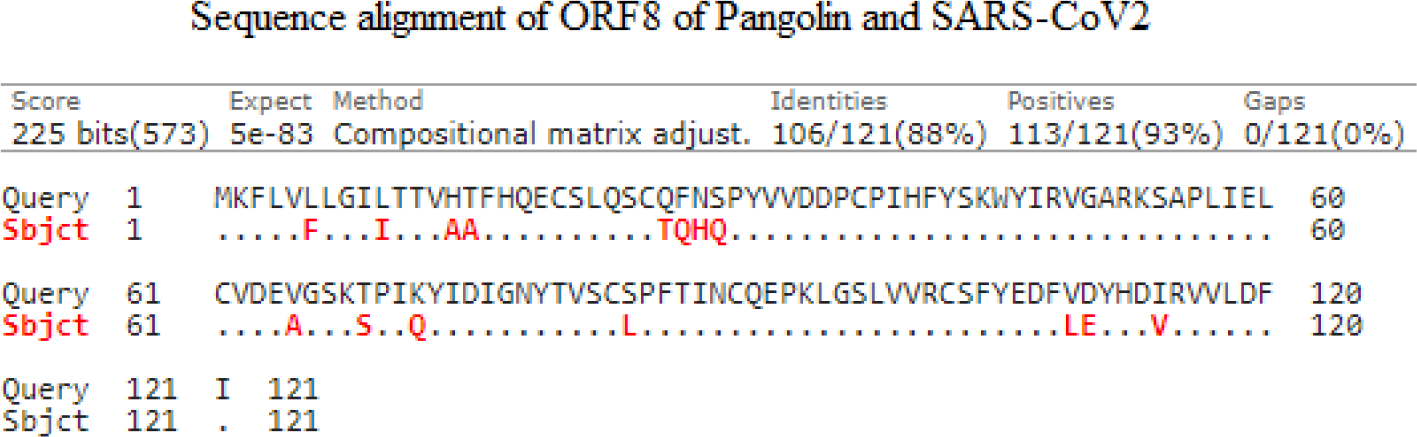
Amino acid sequence alignment of ORF8 proteins of Pangolin-CoV and SARS-CoV-2

We observed a difference of 15 amino acid residues between the Pangolin-CoV and the SARS-CoV-2 ORF8. It was analysed that in both the Bat-CoV RaTG13 and the Pangolin-CoV ORF8 protein the mutations L10I, V65A and S84L are occurred. So, it can be hypothesised that SARS-CoV-2 may have originated from Pangolin-CoV or Bat-CoV RaTG13 ORF8.

Also, the comparison of frequencies of the hydrophobic, hydrophilic and charged amino acids was performed, which are present among the four different ORF8 proteins of SARS-CoV, SARS-CoV-2, Bat RaTG13-CoV and Pangolin-CoV was made. As seen in Table 4, SARS-CoV-2 ORF8, SARS-CoV ORF8ab, Bat-CoV RaTG13 ORF8 and Pangolin-CoV ORF8, are all similar in terms of hydrophobicity and hydrophilicity and it is known that hydrophobicity and hydrophilicity play an important role in protein folding which determines the tertiary structure of the protein and thereby affect the function.

**Table 4:** Comparison of frequencies of hydrophobic, hydrophilic, charged amino acids over four ORF8 proteins

| Properties | SARS-CoV |  | SARS-CoV-2 |  | Bat-CoV RaTG13 | Pangolin-CoV |
| --- | --- | --- | --- | --- | --- | --- |
|  | *ORF8a* | *ORF8b* | *D1 (ORF8)* | *D2 (ORF8)* | *ORF8* | *ORF8* |
| *Hydrophobic* | 17 | 41 | 12 | 46 | 58 | 58 |
| *Hydrophilic* | 22 | 43 | 3 | 60 | 63 | 63 |
| *Positive charged* | 6 | 13 | 1 | 11 | 13 | 14 |
| *Negative charged* | 2 | 3 | 0 | 12 | 13 | 13 |
| *Neutral* | 31 | 68 | 14 | 83 | 95 | 94 |

The ORF8 sequences of SARS-CoV-2, Bat-CoV RaTG13 and Pangolin-CoV have almost the same positive and negative charged amino acids, therefore we can say that probably they have similar kind of electrostatic and hydrophobic interactions, which also contribute to the functionality of the proteins. Again, for the SARS-CoV ORF8ab it was found that the number of positive and negative charged amino acids are closely similar to the SARS-CoV-2 ORF8. Although, the SARS-CoV sequence bears less similarity with the SARS-CoV-2 but they are probably similar in terms of electrostatic and hydrophobic interactions.

Furthermore, we analysed the three ORF8 sequences and checked for their molecular weight, isoelectric point (PI), hydropathy, net charge and extinction coefficient using a peptide property calculator (https://pepcalc.com/) and found that all the properties are almost identical, which shows that the ORF8 protein of Bat-CoV RaTG13, Pangolin-CoV and SARS-CoV-2 are very closely related from the chemical aspects of amino acid residues (Fig. 8).

**Figure 8:**
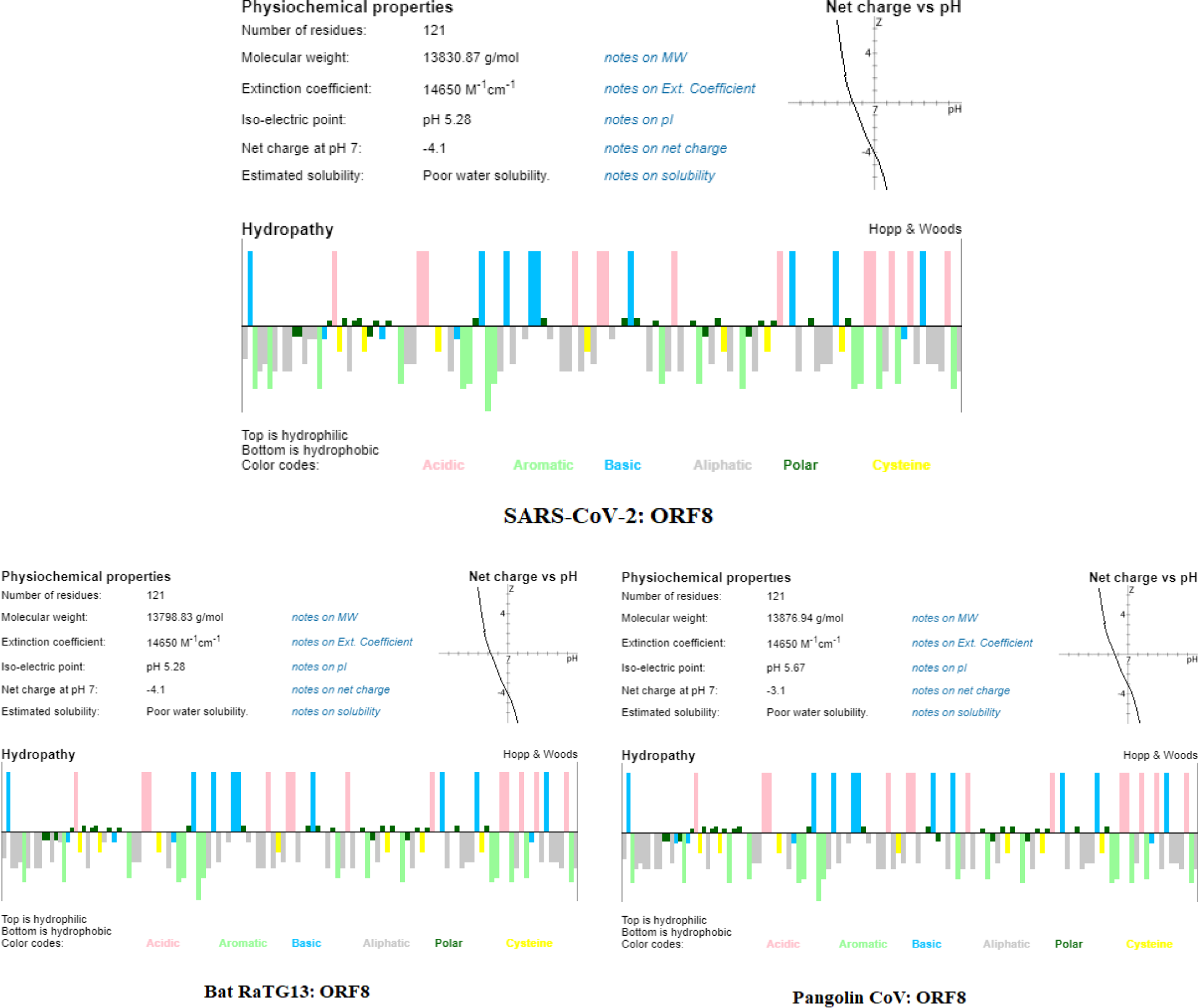
Comparison of peptide properties of ORF8 proteins of SARS-CoV-2 and Bat RaTG13-CoV and Pangolin-CoV

The isoelectric point (PI) and molecular weight of the protein tells us about the biochemical and functional aspects of the protein. Since ORF8 sequences of Bat-CoV RaTG13 and SARS-CoV-2 have the same PI and molecular weights, so they can be grouped under a single functional header. The PI of the Pangolin-CoV ORF8 is higher than SARS-CoV-2 indicating that ORF8 of the Pangolin-CoV is more negatively charged as compared to the SARS-CoV-2 ORF8. Also, since these three proteins have a similar molecular weight, PI, extinction coefficient and nature of hydropathy plot, it would be difficult to differentiate these three proteins by biophysical techniques on the basis of these properties.

The ORF8 of SARS-CoV-2 is significantly different from the ORF8 of Pangolin-CoV and it seems that the ORF8 protein (SARS-CoV-2) is imitating the properties as well the structure of ORF8 of Bat RaTG13-CoV as a blueprint.

The differences between the ORF8 of SARS-CoV-2 and Bat-CoV RaTG13 and Pangolin-CoV can be further demonstrated by the analysis of the per-residue intrinsic disorder predispositions of these proteins. Results of this analysis are shown in Fig. 9A, which illustrates that the intrinsic disorder propensity of the ORF8 from SARS-CoV-2 is closer to that of the ORF8 from Bat-CoV RaTG13 than to the disorder potential of ORF8 from Pangolin-CoV. This is in agreement with the results of other analyses conducted in this study.

**Figure 9:**
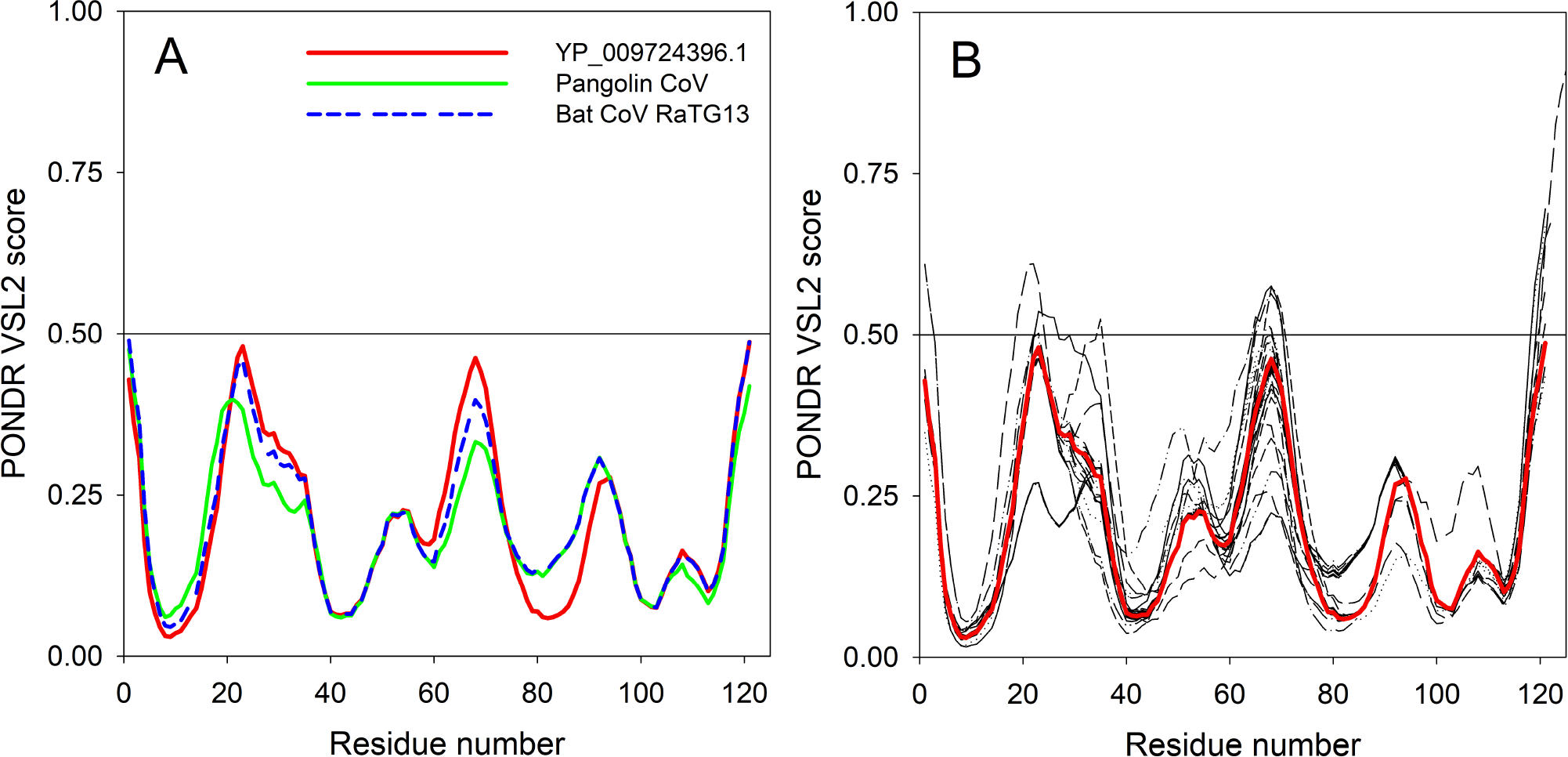
**A.** Comparison of the intrinsic disorder predisposition of the reference ORF8 protein (YP_009724396.1) of the *NC*_045512 SARS-CoV-2 genome from China, Wuhan (bold red curve) with disorder predispositions of ORF8 from the Pangolin-CoV (QIA48620.1) and Bat-CoV RatG13 (QHR63307.1). **B.** Analysis of the intrinsic disorder predisposition of the unique variants of the SARS-CoV-2 ORF8 in comparison with the reference ORF8 protein (YP_009724396.1) of the NC_045512 SARS-CoV-2 genome from China, Wuhan (bold red curve). Analysis is conducted using PONDR-VSL2 algorithm [40], which is one of the more accurate standalone disorder predictors [41, 42, 43]. A disorder threshold is indicated as a thin line (at score = 0.5). Residues/regions with the disorder scores > 0.5 are considered as disordered.

### 3.2. Mutations

Each of the ORF8 amino acid sequences (fasta formatted) are aligned with respect to the ORF8 protein (YP 00972oI396.1) from China-Wuhan using multiple sequence alignment tools (NCBI Blastp suite) and found the mutations and their associated positions were detected accordingly [29]. It is noted that a mutation from an amino acid *A*_1_ to *A*_2_ at a position *p* is denoted by *A*_1_ *pA*_2_ or *A*_1_ (*p*)*A*_2_. The Fig. 10 describes various mutations with their respective locations. The missense mutations are found in the entire ORF8 sequence starting from the amino acid position 3 to 121 and some insertion mutations occurred at the end of the C-terminal. It is found that an amino acid at a fixed position (such as 11, 16, 38, 121 etc.) mutates to multiple amino acids. For example, at position 11 of the reference ORF8 protein, the amino acid threonine (T) maps to isoleucine (I), alanine (A) and lysine (K) in different ORF8 proteins.

**Figure 10:**
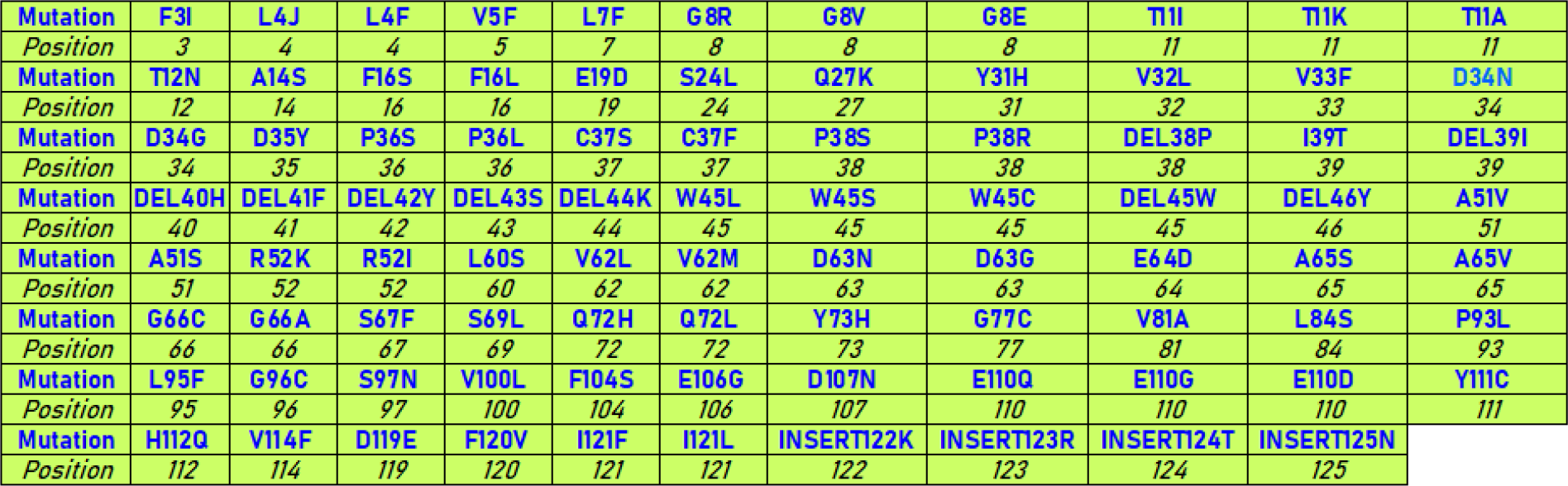
Mutations and their amino acid positions in ORF8 proteins of SARS-CoV-2

Based on observed mutations, it is noticed that amino acids threonine (T) and tryptophan (W) are found to be most vulnerable to mutate to various amino acids. It is noteworthy that the SARS-CoV-2 ORF8 is rapidly undergoing different type of mutations, indicating that it is a highly evolving protein, whereas the Bat-CoV ORF8 is highly conserved (Fig. 6) and the Pangolin-CoV ORF8 is 100% conserved (Fig. 7).

A pie chart presenting the frequency distribution of various mutations is shown in Fig. 11.

**Figure 11:**
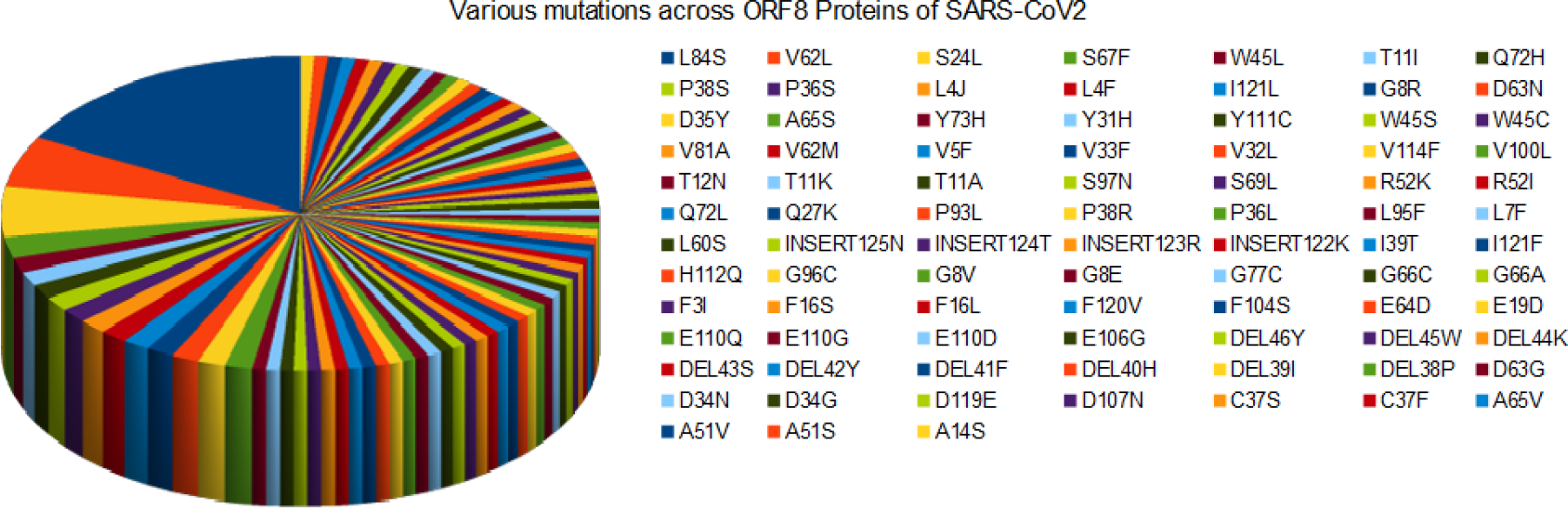
Pie chart of the frequency of distinct mutations of ORF8 proteins of SARS-CoV-2

The N-terminal signal peptide of ORF8 (D1) of SARS-CoV-2 is hydrophobic in nature. We further analysed (Table 5) the mutations and observed that hydrophobic to hydrophobic mutations are dominating, indicating that hydrophobicity of the domain is maintained and thus we can postulate that there is probably no functional change in the hydrophobic N-terminal signal peptide.

**Table 5:** Change of amino acid residues on the basis of hydrophobic and hydrophilic properties

| Properties | Number of residues changed the properties |  |
| --- | --- | --- |
|  | D1 | D2 |
| *Hydrophobic to Hydrophobic* | 6 | 14 |
| *Hydrophobic to Hydrophilic* | 3 | 19 |
| *Hydrophilic to Hydrophobic* | 2 | 9 |
| *Hydrophilic to Hydrophilic* | 2 | 19 |

Further, it was found that there was a change from hydrophilicity to hydrophobicity in two positions, thus enhancing the hydrophobic nature of D1. Although, hydrophobic to hydrophilic and hydrophilic to hydrophilic mutations were also observed, they were insignificant when compared to hydrophobicity changes as hydrophobic mutations were observed in eight positions, whereas hydrophilic mutations were only present in five positions. The ORF8 chain protein (D2) was demonstrated to be a region enriched hydrophilic mutations. The hydrophilic mutations were found in about thirty-eight positions. In contrast, there were only twenty-three hydrophobic mutations in D2 thus, enhancing the hydrophilicity and contributing to the soluble nature of the protein.

Distinct non-synonymous mutations and the associated frequency of mutations, predicted effect (using Meta-SNP) as well as the predicted change of structural stability (using I-MUTANT) due to mutation(s) are presented in Table 6. The most frequent mutation in the ORF8 proteins turned out to be L84S (hydrophobic (L) to non-charged hydrophilic (S)) which is a clade (S) determining mutation with frequency 23 [44]. It is observed from Table 6 that the mutation L84S decreases structural stability and consequently changes the functions of ORF8. Note that the ORF8 sequence QLJ93922.1 (USA) possesses consecutive (38-46 aa) deletion mutations. Also the other sequence QKI36860.1 (China: Guangzhou) accommodates four insertion mutations at the end of the C terminal (122-125) along with two other mutations S84L and D119E.

**Table 6:**
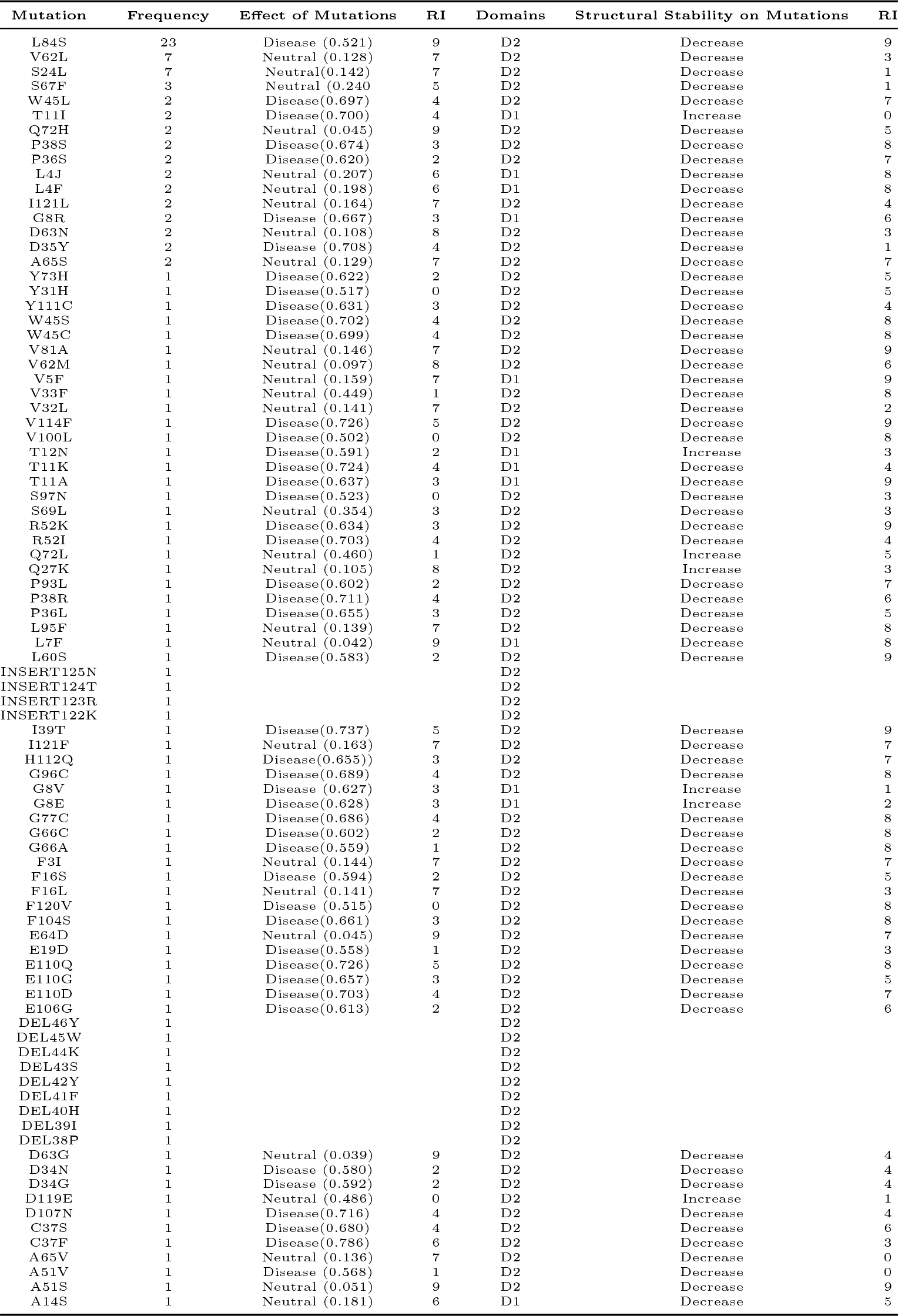
Mutations across ORF8 proteins of SARS-CoV-2 and their predicted effects

| Mutation | Frequency | Effect of Mutations | RI | Domains | Structural Stability on Mutations | RI |
| --- | --- | --- | --- | --- | --- | --- |
| L84S | 23 | Disease (0.521) | 9 | D2 | Decrease | 9 |
| V62L | 7 | Neutral (0.128) | 7 | D2 | Decrease | 3 |
| S24L | 7 | Neutral(0.142) | 7 | D2 | Decrease | 1 |
| S67F | 3 | Neutral (0.240) | 5 | D2 | Decrease | 1 |
| W45L | 2 | Disease(0.697) | 4 | D2 | Decrease | 7 |
| T11I | 2 | Disease(0.700) | 4 | D1 | Increase | 0 |
| Q72H | 2 | Neutral (0.045) | 9 | D2 | Decrease | 5 |
| P38S | 2 | Disease(0.674) | 3 | D2 | Decrease | 8 |
| P36S | 2 | Disease(0.620) | 2 | D2 | Decrease | 7 |
| L4J | 2 | Neutral (0.207) | 6 | D1 | Decrease | 8 |
| L4F | 2 | Neutral (0.198) | 6 | D1 | Decrease | 8 |
| I121L | 2 | Neutral (0.164) | 7 | D2 | Decrease | 4 |
| G8R | 2 | Disease (0.667) | 3 | D1 | Decrease | 6 |
| D63N | 2 | Neutral (0.108) | 8 | D2 | Decrease | 3 |
| D35Y | 2 | Disease (0.708) | 4 | D2 | Decrease | 1 |
| A65S | 2 | Neutral (0.129) | 7 | D2 | Decrease | 7 |
| Y73H | 1 | Disease(0.622) | 2 | D2 | Decrease | 5 |
| Y31H | 1 | Disease(0.517) | 0 | D2 | Decrease | 5 |
| Y111C | 1 | Disease(0.631) | 3 | D2 | Decrease | 4 |
| W45S | 1 | Disease(0.702) | 4 | D2 | Decrease | 8 |
| W45C | 1 | Disease(0.699) | 4 | D2 | Decrease | 8 |
| V81A | 1 | Neutral (0.146) | 7 | D2 | Decrease | 9 |
| V62M | 1 | Neutral (0.097) | 8 | D2 | Decrease | 6 |
| V5F | 1 | Neutral (0.159) | 7 | D1 | Decrease | 9 |
| V33F | 1 | Neutral (0.449) | 1 | D2 | Decrease | 8 |
| V32L | 1 | Neutral (0.141) | 7 | D2 | Decrease | 2 |
| V114F | 1 | Disease(0.726) | 5 | D2 | Decrease | 9 |
| V100L | 1 | Disease(0.502) | 0 | D2 | Decrease | 8 |
| T12N | 1 | Disease(0.591) | 2 | D1 | Increase | 3 |
| T11K | 1 | Disease(0.724) | 4 | D1 | Decrease | 4 |
| T11A | 1 | Disease(0.637) | 3 | D1 | Decrease | 9 |
| S97N | 1 | Disease(0.523) | 0 | D2 | Decrease | 3 |
| S69L | 1 | Neutral (0.354) | 3 | D2 | Decrease | 3 |
| R52K | 1 | Disease(0.634) | 3 | D2 | Decrease | 9 |
| R52I | 1 | Disease(0.703) | 4 | D2 | Decrease | 4 |
| Q72L | 1 | Neutral (0.460) | 1 | D2 | Increase | 5 |
| Q27K | 1 | Neutral (0.105) | 8 | D2 | Increase | 3 |
| P93L | 1 | Disease(0.602) | 2 | D2 | Decrease | 7 |
| P38R | 1 | Disease(0.711) | 4 | D2 | Decrease | 6 |
| P36L | 1 | Disease(0.655) | 3 | D2 | Decrease | 5 |
| L95F | 1 | Neutral (0.139) | 7 | D2 | Decrease | 8 |
| L7F | 1 | Neutral (0.042) | 9 | D1 | Decrease | 8 |
| L60S | 1 | Disease(0.583) | 2 | D2 | Decrease | 9 |
| INSERT125N | 1 |  |  | D2 |  |  |
| INSERT124T | 1 |  |  | D2 |  |  |
| INSERT123R | 1 |  |  | D2 |  |  |
| INSERT122K | 1 |  |  | D2 |  |  |
| I39T | 1 | Disease(0.737) | 5 | D2 | Decrease | 9 |
| I121F | 1 | Neutral (0.163) | 7 | D2 | Decrease | 7 |
| H112Q | 1 | Disease(0.655)) | 3 | D2 | Decrease | 7 |
| G96C | 1 | Disease(0.689) | 4 | D2 | Decrease | 8 |
| G8V | 1 | Disease (0.627) | 3 | D1 | Increase | 1 |
| G8E | 1 | Disease(0.628) | 3 | D1 | Increase | 2 |
| G77C | 1 | Disease(0.686) | 4 | D2 | Decrease | 8 |
| G66C | 1 | Disease(0.602) | 2 | D2 | Decrease | 8 |
| G66A | 1 | Disease(0.559) | 1 | D2 | Decrease | 8 |
| F3I | 1 | Neutral (0.144) | 7 | D2 | Decrease | 7 |
| F16S | 1 | Disease (0.594) | 2 | D2 | Decrease | 5 |
| F16L | 1 | Neutral (0.141) | 7 | D2 | Decrease | 3 |
| F120V | 1 | Disease (0.515) | 0 | D2 | Decrease | 8 |
| F104S | 1 | Disease(0.661) | 3 | D2 | Decrease | 8 |
| E64D | 1 | Neutral (0.045) | 9 | D2 | Decrease | 7 |
| E19D | 1 | Disease(0.558) | 1 | D2 | Decrease | 3 |
| E110Q | 1 | Disease(0.726) | 5 | D2 | Decrease | 8 |
| E110G | 1 | Disease(0.657) | 3 | D2 | Decrease | 5 |
| E110D | 1 | Disease(0.703) | 4 | D2 | Decrease | 7 |
| E106G | 1 | Disease(0.613) | 2 | D2 | Decrease | 6 |
| DEL46Y | 1 |  |  | D2 |  |  |
| DEL45W | 1 |  |  | D2 |  |  |
| DEL44K | 1 |  |  | D2 |  |  |
| DEL43S | 1 |  |  | D2 |  |  |
| DEL42Y | 1 |  |  | D2 |  |  |
| DEL41F | 1 |  |  | D2 |  |  |
| DEL40H | 1 |  |  | D2 |  |  |
| DEL39I | 1 |  |  | D2 |  |  |
| DEL38P | 1 |  |  | D2 |  |  |
| D63G | 1 | Neutral (0.039) | 9 | D2 | Decrease | 4 |
| D34N | 1 | Disease (0.580) | 2 | D2 | Decrease | 4 |
| D34G | 1 | Disease (0.592) | 2 | D2 | Decrease | 4 |
| D119E | 1 | Neutral (0.486) | 0 | D2 | Increase | 1 |
| D107N | 1 | Disease(0.716) | 4 | D2 | Decrease | 4 |
| C37S | 1 | Disease(0.680) | 4 | D2 | Decrease | 6 |
| C37F | 1 | Disease(0.786) | 6 | D2 | Decrease | 3 |
| A65V | 1 | Neutral (0.136) | 7 | D2 | Decrease | 0 |
| A51V | 1 | Disease (0.568) | 1 | D2 | Decrease | 0 |
| A51S | 1 | Neutral (0.051) | 9 | D2 | Decrease | 9 |
| A14S | 1 | Neutral (0.181) | 6 | D1 | Decrease | 5 |

Based on predicted effects and change of structural stability, mutations are grouped into four classes as shown in Table 7

**Table 7:** Classification of the missense mutations based on predicted effects as well as change of structural stability

| *D-D | D-D | D-D | *D-I | *N-D | N-D | *N-I |
| --- | --- | --- | --- | --- | --- | --- |
| L84S | T11A | G66A | *T11I* | V62L | V33F | *Q72L* |
| P38S | S97N | F16S | *T12N* | S24L | V32L | *Q27K* |
| P36S | R52K | F120V | *G8V* | S67F | S69L | *D119E* |
| G8R | R52I | F104S | *G8E* | Q72H | L95F |  |
| D35Y | P93L | E19D |  | L4J | L7F |  |
| Y73H | P38R | E110Q |  | L4F | I121F |  |
| Y31H | P36L | E110G |  | I121L | F3I |  |
| Y111C | L60S | E110D |  | D63N | F16L |  |
| W45S | I39T | E106G |  | A65S | E64D |  |
| W45C | H112Q | D34N |  | V81A | D63G |  |
| V114F | G96C | D34G |  | V62M | A65V |  |
| V100L | G77C | D107N |  | V5F | A51S |  |
| T11K | G66C | C37S |  |  | A14S |  |
|  | W45L | C37F |  |  |  |  |
|  |  | A51V |  |  |  |  |

\*D-D: Disease-Decreasing; \*N-D: Neutral-Decreasing; \*D-I:Disease-Increasing and \*N-I: Neutral-Increasing.

Based on predicted effect and change of stability we classified the mutations into four types (Table 7):

1. **Disease-Decreasing**: This class has higher frequency of mutations showing that most of the disease mutations are decreasing the stability of the protein and most of them occurred in D2.
2. **Neutral-Decreasing**: Although the mutations are of neutral type and supposedly are not harmful for the host, they cause the stability of protein structure to decrease.
3. **Disease-Increasing**: These mutations lie in the D1 and increase the stability of the protein, making the hydrophobic N-terminal more stable and thereby enhancing the localisation to ER more efficient.
4. **Neutral-Increasing**: The frequency of mutations are very low in this class, although the mutations are neutral but they increase the stability of the protein effectively and they all occur in the D2 domain.

In Table 8, the list of unique ORF8 protein IDs and their associated mutations with domain(s) and the predicted effects and changes of structural stability are presented.

**Table 8:** ORF8 protein IDs and corresponding mutations, predicted effects

| Protein ID | Mutation(s) | Predicted effect of mutations | RI | Predicted effect on stability | RI | Domain(s) |
| --- | --- | --- | --- | --- | --- | --- |
| QMI92505.1 | L4F | Neutral (0.198) | 6 | Decreases | 8 | D1 |
| QMT48896.1 | L4F, D63N | Neutral (0.198), Neutral (0.108) | 6, 8 | Decreases, Decrease | 8,3 | D1, D2 |
| QKU31272.1 | L4I | Neutral (0.207) | 6 | Decreases | 8 | D1 |
| QLG99743.1 | L4I | Neutral (0.207) | 6 | Decreases | 8 | D1 |
| QMT28672.1 | V5F, L84S | Neutral (0.159), Disease (0.521) | 7, 9 | Decreases, Decrease | 9,9 | D1, D2 |
| QMS51860.1 | L7F | Neutral (0.042) | 9 | Decrease | 8 | D1 |
| QJY40584.1 | G8E | Disease(0.628) | 3 | Increase | 2 | D1 |
| QKV07730.1 | T11A, L84S | Disease(0.637), Disease (0.521) | 3, 9 | Decrease, Decrease | 9,9 | D1,D2 |
| QMU94963.1 | T11I | Disease(0.700) | 4 | Increase | 0 | D1 |
| QMS53022.1 | T11I, L84S | Disease(0.700), Disease (0.521) | 4, 9 | Increase, Decrease | 0,9 | D1,D2 |
| QKE49215.1 | T11K | Disease(0.724) | 4 | Decrease | 4 | D1 |
| QMU91718.1 | T12N | Disease(0.591) | 2 | Increase | 3 | D1 |
| QMT50804.1 | E19D, L84S | Disease(0.558), Disease (0.521) | 1, 9 | Decrease, Decrease | 3,9 | D2,D2 |
| QJS56890.1 | S24L, W45S | Neutral(0.142), Disease(0.702) | 7, 4 | Decrease, Decrease | 1,8 | D2,D2 |
| QLH58821.1 | S24L, V62L | Neutral(0.142), Neutral (0.128) | 7, 7 | Decrease, Decrease | 1,3 | D2, D2 |
| QLY90504.1 | S24L, S67F | Neutral(0.142), Neutral (0.240) | 7, 5 | Decrease, Decrease | 1,1 | D2,D2 |
| QMU91370.1 | S24L, S69L | Neutral(0.142), Neutral (0.354) | 7, 3 | Decrease, Decrease | 1,3 | D2,D2 |
| QMV29747.1 | S24L | Neutral(0.142) | 7 | Decrease | 1 | D2 |
| QJR91288.1 | S24L | Neutral(0.142) | 7 | Decrease | 1 | D2 |
| QLH58953.1 | S24L, P38S | Neutral(0.142), Disease(0.674) | 7, 3 | Decrease, Decrease | 1,8 | D2,D2 |
| QMJ00652.1 | Q27K | Neutral (0.105) | 8 | Increase | 3 | D2 |
| QLC47867.1 | Y31H | Disease(0.517) | 0 | Decrease | 5 | D2 |
| QMU91598.1 | V32L | Neutral (0.141) | 7 | Decrease | 2 | D2 |
| QLM05830.1 | V33F | Neutral (0.449) | 1 | Decrease | 8 | D2 |
| QKQ63407.1 | P36L | Disease(0.655) | 3 | Decrease | 5 | D2 |
| QMT55731.1 | W45C | Disease(0.699) | 4 | Decrease | 8 | D2 |
| QLL35457.1 | W45L | Disease(0.697) | 4 | Decrease | 7 | D2 |
| QKU37052.1 | W45L | Disease(0.697), Neutral (0.479) | 4, 0 | Decrease, Decrease | 7,9 | D2, D2 |
| QMT49652.1 | R52I | Disease(0.703) | 4 | Decrease | 4 | D2 |
| QLK98030.1 | R52K | Disease(0.634) | 3 | Decrease | 9 | D2 |
| QMU93583.1 | V62L | Neutral (0.128) | 7 | Decrease | 3 | D2 |
| QMU91550.1 | V62L, L84S | Neutral (0.128), Disease (0.521) | 7, 9 | Decrease, Decrease | 3, 9 | D2,D2 |
| QJR88936.1 | V62L, L84S | Neutral (0.128), Disease (0.521) | 7, 9 | Decrease, Decrease | 3, 9 | D2,D2 |
| QIZ15700.1 | V62L, L84S | Neutral (0.128), Disease (0.521) | 7, 9 | Decrease, Decrease | 3, 9 | D2,D2 |
| QMS95048.1 | V62M | Neutral (0.097) | 8 | Decrease | 6 | D2 |
| QLF78328.1 | E64D | Neutral (0.045) | 9 | Decrease | 7 | D2 |
| QJS57274.1 | G66A | Disease(0.559) | 1 | Decrease | 8 | D2 |
| QKQ29929.1 | G66C | Disease(0.602) | 2 | Decrease | 8 | D2 |
| QMT56283.1 | S67F | Neutral (0.240) | 5 | Decrease | 1 | D2 |
| QKV06506.1 | S67F, L84S | Neutral (0.240), Disease (0.521) | 5, 9 | Decrease, Decrease | 1, 9 | D2,D2 |
| QMT91163.1 | Q72H | Neutral (0.045) | 9 | Decrease | 5 | D2 |
| QKV40062.1 | Q72H, L84S | Neutral (0.045), Disease (0.521) | 9, 9 | Decrease, Decrease | 5,9 | D2,D2 |
| QLA47566.1 | Q72L | Neutral (0.460) | 1 | Increase | 5 | D2 |
| QMI96758.1 | Y73H | Disease(0.622) | 2 | Decrease | 5 | D2 |
| QKV38912.1 | G77C | Disease(0.686) | 4 | Decrease | 8 | D2 |
| QMI91809.1 | V81A | Neutral (0.146) | 7 | Decrease | 9 | D2 |
| QMT95795.1 | P93L | Disease(0.602) | 2 | Decrease | 7 | D2 |
| QMU84881.1 | L95F | Neutral (0.139) | 7 | Decrease | 8 | D2 |
| QMI92229.1 | G96C | Disease(0.689) | 4 | Decrease | 8 | D2 |
| QMT55779.1 | S97N | Disease(0.523) | 0 | Decrease | 3 | D2 |
| QJC19630.1 | V100L | Disease(0.502) | 0 | Decrease | 8 | D2 |
| QMU84893.1 | E110D | Disease(0.703) | 4 | Decrease | 7 | D2 |
| QMJ01228.1 | Y111C | Disease(0.631) | 3 | Decrease | 4 | D2 |
| QJR88180.1 | V114F | Disease(0.726) | 5 | Decrease | 9 | D2 |
| QJT72369.1 | I121F | Neutral (0.163) | 7 | Decrease | 7 | D2 |
| QJT43615.1 | I121L | Neutral (0.164) | 7 | Decrease | 4 | D2 |
| QMJ01216.1 | F120V, I121L | Disease (0.515), Neutral (0.164) | 0, 7 | Decrease, Decrease | 8,4 | D2,D2 |
| QLJ58008.1 | E110G | Disease(0.657) | 3 | Decrease | 5 | D2 |
| QJW28311.1 | D107N | Disease(0.716) | 4 | Decrease | 4 | D2 |
| QJS53933.1 | E106G | Disease(0.613) | 2 | Decrease | 6 | D2 |
| QMU91358.1 | L84S | Disease (0.521) | 9 | Decrease | 9 | D2 |
| QJR90724.1 | L84S | Disease (0.521) | 9 | Decrease | 9 | D2 |
| QKI36860.1 (*) | L84S, D119E | Disease (0.521), Neutral (0.486) | 9 | Decrease, Increase | 9,1 | D2,D2 |
| QJD48694.1 | L84S, H112Q | Disease (0.521),Disease(0.655)) | 9, 3 | Decrease, Decrease | 9,7 | D2,D2 |
| QMS54342.1 | L84S, E110Q | Disease (0.521), Disease(0.726) | 9, 5 | Decrease, Decrease | 9, 8 | D2,D2 |
| QMT96539.1 | L84S, F104S | Disease (0.521), Disease(0.661) | 9, 3 | Decrease, Decrease | 9, 9 | D2,D2 |
| QMT51608.1 | A65S | Neutral (0.129) | 7 | Decrease | 7 | D2 |
| QLH01196.1 | A65S, L84S | Neutral (0.129), Disease (0.521) | 7, 9 | Decrease, Decrease | 7,9 | D2,D2 |
| QMU91298.1 | A65V | Neutral (0.136) | 7 | Decrease | 0 | D2 |
| QKU32832.1 | L84S | Disease (0.521) | 9 | Decrease | 9 | D2 |
| QMU91334.1 | D63G, L84S | Neutral (0.039), Disease (0.521) | 9, 9 | Decrease, Decrease | 4, 9 | D2,D2 |
| QMT50180.1 | D63N | Neutral (0.108) | 8 | Decrease | 3 | D2 |
| QKC05159.1 | L84S | Disease (0.521) | 9 | Decrease | 9 | D2 |
| QMU94207.1 | L60S | Disease(0.583) | 2 | Decrease | 9 | D2 |
| QJX70307.1 | A51S | Neutral (0.051) | 9 | Decrease | 9 | D2 |
| QMT54388.1 | A51V | Disease (0.568) | 1 | Decrease | 0 | D2 |
| QLH01532.1 | I39T | Disease(0.737) | 5 | Decrease | 9 | D2 |
| QLF97742.1 | P38R | Disease(0.711) | 4 | Decrease | 6 | D2 |
| QJZ28306.1 | P38S | Disease(0.674) | 3 | Decrease | 8 | D2 |
| QMT56763.1 | C37F | Disease(0.786) | 6 | Decrease | 3 | D2 |
| QMT50144.1 | C37S | Disease(0.680) | 4 | Decrease | 6 | D2 |
| QKG86865.1 | P36S, V62L, L84S | Disease(0.620),Neutral (0.128),Neutral (0.479) | 2, 7, 0 | Decrease, Decrease, Decrease | 7,3,9 | D2, D2, D2 |
| QJR93136.1 | P36S | Disease (0.620) | 2 | Decrease | 7 | D2 |
| QJR88780.1 | L84S | Disease (0.521) | 9 | Decrease | 9 | D2 |
| QLJ84632.1 | D35Y | Disease (0.708) | 4 | Decrease | 1 | D2 |
| QLB39040.1 | D34G | Disease (0.592) | 2 | Decrease | 4 | D2 |
| QMU25342.1 | D34N | Disease (0.580) | 2 | Decrease | 4 | D2 |
| QLH57924.1 | F16L, V62L, L84S | Neutral (0.141), Neutral (0.128), Disease (0.521) | 7,7,9 | Decrease, Decrease, Decrease | 3,3,9 | D2,D2,D2 |
| QKV39247.1 | F16S | Disease (0.594) | 2 | Decrease | 5 | D2 |
| QLQ87643.1 | A14S | Neutral (0.181) | 6 | Decrease | 5 | D1 |
| QMT96239.1 | G8R | Disease (0.667) | 3 | Decrease | 6 | D1 |
| QMU92030.1 | G8R, D35Y | Disease (0.667), Disease (0.708) | 3, 4 | Decrease, Decrease | 6,1 | D1,D2 |
| QJR93160.1 | G8V | Disease (0.627) | 3 | Increase | 1 | D1 |
| QLQ87643.1 | A14S | Neutral (0.181) | 0 | Decrease | 7 | D1 |

Further, based on the three different types of mutations viz. neutral, disease and mix of neutral & disease, all the ORF8 proteins are classified into three groups which are adumbrated in Table 9. Also the corresponding pie chart is given in Fig. 12.

**Figure 12:**
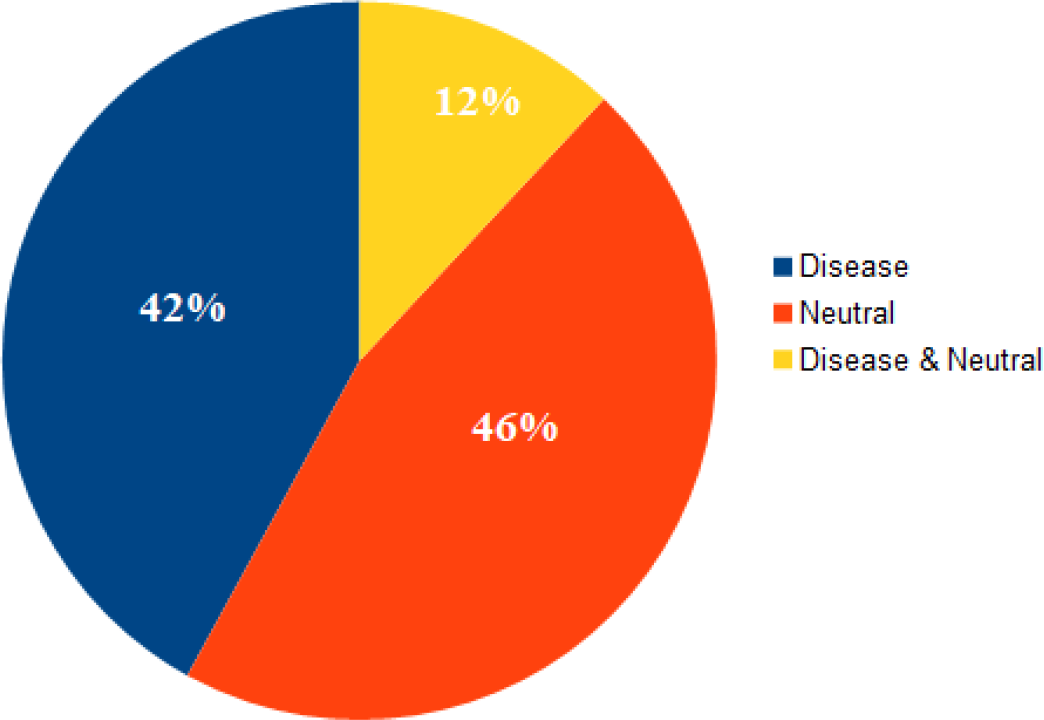
Pie chart of three types of mutations across various ORF8 proteins of SARS-CoV-2

**Table 9:** List of the three type of mutations possessed by 95 distinct ORF8 proteins

| Disease Type |  | Neutral Type |  | Disease & Neutral Type |  |
| --- | --- | --- | --- | --- | --- |
| Accession | Geo.Location | Accession | Geo.Location | Accession | Geo.Location |
| QMU84893 | Bangladesh | QMV29747 | USA: SC | QMT50804 | USA: Washington,Spokane County |
| QMU91718 | USA: FL | QMU84881 | Bangladesh | QMT96539 | USA: FL |
| QMU92030 | USA: FL | QMU91298 | USA: FL | QMS53022 | USA: Washington,Yakima County |
| QMU94207 | USA: Illinois, Cook county | QMU91334 | USA: FL | QMS54342 | USA: Washington,King County |
| QMU94963 | Bangladesh | QMU91358 | USA: FL | QLJ58008 | USA |
| QMU25342 | USA: MD | QMU91370 | USA: FL | QLH58953 | USA: FL |
| QMT49652 | USA: Washington,Yakima County | QMU91550 | USA: FL | QKU37052 | Saudi Arabia: Jeddah |
| QMT50144 | USA: Washington,Yakima County | QMU91598 | USA: FL | QKV07730 | USA: Washington,Asotin County |
| QMT54388 | USA: Washington,Yakima County | QMU93583 | USA: Wisconsin, Dane county | QKG86865 | USA: Massachusetts |
| QMT55731 | USA: Washington,Yakima County | QMT48896 | USA: Washington,Yakima County | QJS56890 | USA: ID |
| QMT55779 | USA: Washington,Yakima County | QMT50180 | USA: Washington,Yakima County | QJD48694 | USA: WA |
| QMT56763 | USA: Washington,Yakima County | QMT51608 | USA: Washington,Yakima County |  |  |
| QMT95795 | USA: Washington,Spokane County | QMT56283 | USA: Washington,Yakima County |  |  |
| QMT96239 | USA: FL | QMT91163 | USA: Washington,Yakima County |  |  |
| QMI92229 | USA: California | QMT28672 | USA |  |  |
| QMI96758 | USA: San Diego, California | QMS95048 | Iran |  |  |
| QMJ01228 | USA: San Diego, California | QMS51860 | USA: Washington,Yakima County |  |  |
| QLL35457 | USA: FL | QMI91809 | USA: San Diego, California |  |  |
| QLK98030 | USA: IN | QMI92505 | USA: San Diego, California |  |  |
| QLJ84632 | France | QMJ00652 | USA: San Diego, California |  |  |
| QLH01532 | USA: CA | QMJ01216 | USA: San Diego, California |  |  |
| QLF97742 | Bangladesh | QLY90504 | USA |  |  |
| QLC47867 | USA: FL | QLQ87643 | India: Surat |  |  |
| QLB39040 | USA | QLM05830 | USA: Wisconsin, Dane county |  |  |
| QKV38912 | USA: Washington, Snohomish County | QLH57924 | USA: FL |  |  |
| QKV39247 | USA: Washington, Snohomish County | QLH58821 | USA: FL |  |  |
| QKQ29929 | Bangladesh: Dhaka | QLG99743 | USA: CA |  |  |
| QKQ63407 | USA: Virginia | QLH01196 | USA: CA |  |  |
| QKE49215 | USA: CA | QLF78328 | Poland |  |  |
| QJZ28306 | Poland | QLA47566 | USA: Virginia |  |  |
| QJY40584 | India: Una | QKV40062 | USA: Washington, Yakima County |  |  |
| QJW28311 | USA: VA | QKU31272 | USA: CA |  |  |
| QJT72369 | France | QKU32832 | USA: CA |  |  |
| QJS53933 | Greece: Athens | QKV06506 | USA: Washington,Mason County |  |  |
| QJS57274 | USA: CT | QKI36860 | China: Guangzhou |  |  |
| QJR88180 | Australia: Victoria | QKC05159 | USA |  |  |
| QJR93136 | Australia: Victoria | QJX70307 | USA |  |  |
| QJR93160 | Australia: Victoria | QJT43615 | India: Ahmedabad |  |  |
| QJC19630 | USA: CT | QJR88780 | Australia: Victoria |  |  |
|  |  | QJR88936 | Australia: Victoria |  |  |
|  |  | QJR90724 | Australia: Victoria |  |  |
|  |  | QJR91288 | Australia: Victoria |  |  |
|  |  | QIZ15700 | USA: GA |  |  |

From Table 9, it was concluded that the majority of mutations examined in the distinct variants of the ORF8 proteins of SARS-CoV-2 are turned up to be neutral, while 42% of the mutations become disease-causing as predicted. Furthermore, based on the presence of the most important mutations L84S and S24L in unique variants of the ORF8 (SARS-CoV-2), two strains are proposed, although one of which (L84S) is already defined in the literature [44]. In Table 10 and 11, the list of ORF8 protein IDs with associated details are presented.

**Table 10:** ORF8 proteins containing the strain determining mutation L84S

| Accession | Geo_Location | Collection_Date | Accession | Geo_Location | Collection_Date |
| --- | --- | --- | --- | --- | --- |
| QKI36860 | China: Guangzhou | 2020-02-25 | QMT96539 | USA: FL | 2020-03-21 |
| QIZ15700 | USA: GA | 2020-03 | QMU91550 | USA: FL | 2020-03-24 |
| QKC05159 | USA | 2020-03-07 | QJR90724 | Australia: Victoria | 2020-03-25 |
| QJD48694 | USA: WA | 2020-03-11 | QKG86865 | USA: Massachusetts | 2020-03-26 |
| QMU91334 | USA: FL | 2020-03-13 | QKV07730 | USA: Washington,Asotin County | 2020-04-01 |
| QMU91358 | USA: FL | 2020-03-13 | QKV06506 | USA: Washington,Mason County | 2020-04-02 |
| QMT28672 | USA | 2020-03-13 | QLH01196 | USA: CA | 2020-04-11 |
| QKU37052 | Saudi Arabia: Jeddah | 2020-03-15 | QLH57924 | USA: FL | 2020-04-15 |
| QKU32832 | USA: CA | 2020-03-18 | QKV40062 | USA: Washington, Yakima County | 2020-05-01 |
| QJR88780 | Australia: Victoria | 2020-03-20 | QMS54342 | USA: Washington,King County | 2020-05-04 |
| QJR88936 | Australia: Victoria | 2020-03-20 | QMS53022 | USA: Washington,Yakima County | 2020-05-16 |
|  |  |  | QMT50804 | USA: Washington,Spokane County | 2020-06-09 |

**Table 11:**
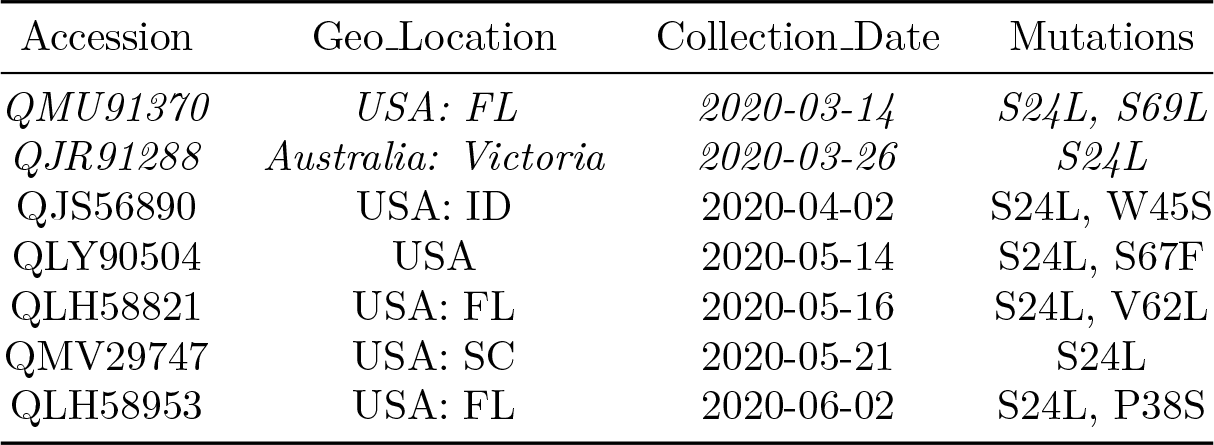
ORF8 proteins containing the strain determining mutation S24L

Note that, other than these two strains, there are other sixty-four other ORF8 sequences, which do not possess any of these two strain-determining mutations. This clarifies that the ORF8 protein is certainly one of the fundamental proteins which directs the pathogenicity of a variety of strains of SARS-CoV-2.

#### 3.2.1. Remarks Based on Mutations over ORF8 Proteins of SARS-CoV-2, Bat-CoV and Pangolin-CoV

To study the evolution of mutations and to observe the relationships among three ORF8 proteins, we compared SARS-CoV-2 ORF8 with that of Bat-CoV RaTG13 and Pangolin-CoV ORF8. The detailed analysis of all mutations is presented in Table 12.

**Table 12:**
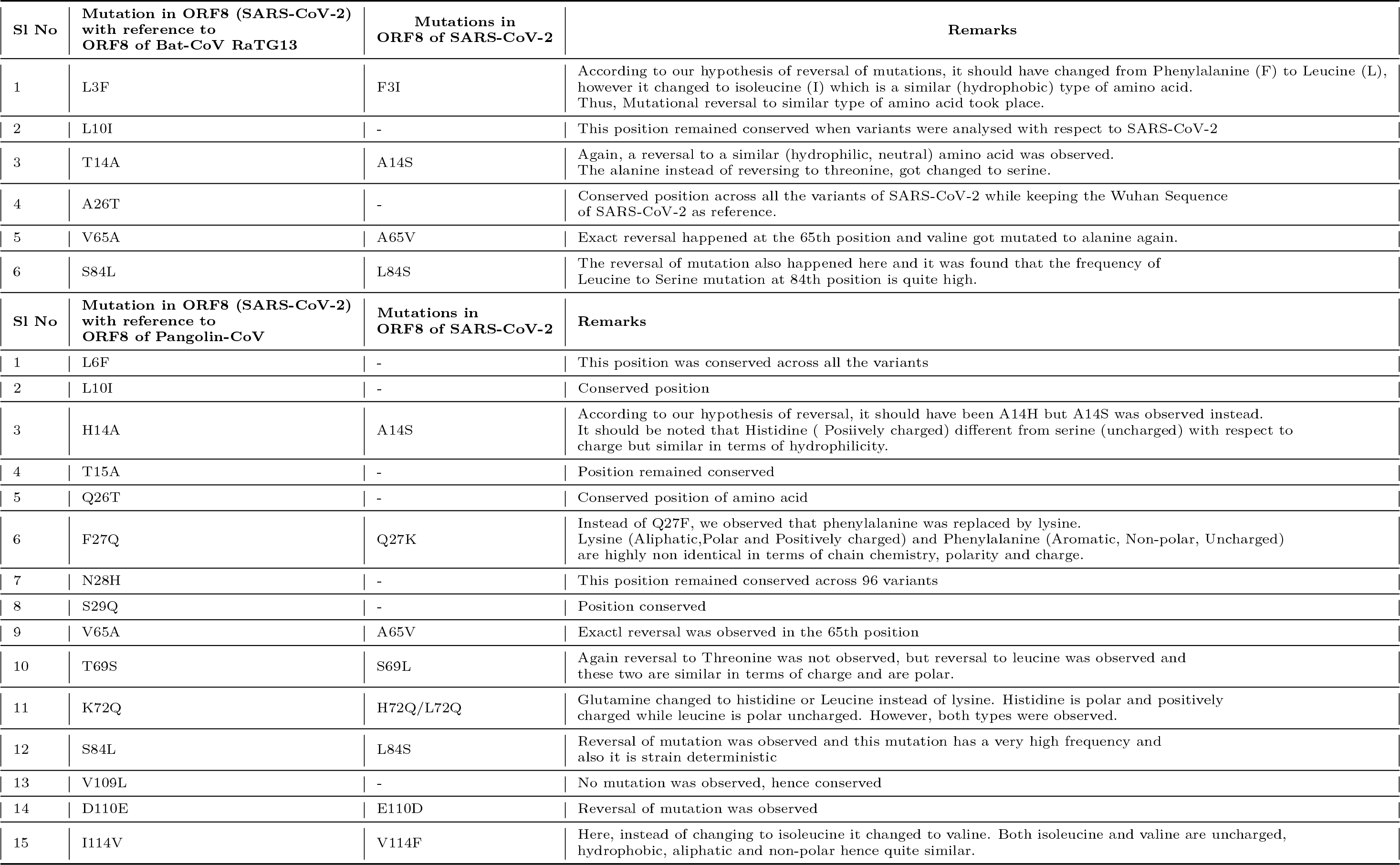
Observations on mutations in ORF8 with reference to ORF8 of Pangolin-CoV, Bat-CoVs

Based on Table 12, it can be suggested that reverse mutations will lead to the same sequence for the SARS-CoV-2 ORF8 protein will have the same sequence as that of Bat-CoV RaTG13 and Pangolin-CoV ORF8 proteins in the near future. We can also conclude that SARS-CoV-2 is showing genetically reverse engineering when compared with Bat-CoV RaTG13 and Pangolin-CoV ORF8.

Fig. 9 (**B**) represents intrinsic disorder profiles for 96 variants of the ORF8 protein from different isolates of SARS-CoV-2. The intrinsic disorder predispositions can vary significantly, especially within the highly and moderately flexible regions. Although many mutations are disorder-silent, some of the mutations increase the local disorder propensity, whereas other cause a noticeable decrease in the disorder predisposition. For example, local disorder predisposition in the vicinity of residue 20 was increased in the variants QKV39247.1 and QLC47867.1. Variants QMT50144.1 and QLH01532.1 had a prominent new peak in the vicinity of residue 30, where the vast majority of other variants have a shoulder. Although variant QJS56890.1 also has a prominent peak in the vicinity of residue 30 and has one of the highest peaks in the vicinity of residue 50, the intensity of its peak in the vicinity of residue 20 is noticeably decreased. Variant QMT96539.1 showed higher disorder propensity in the vicinity of residue 110. Variants QMT49652.1 and QMT54388.1 have the lowest disorder predisposition in the vicinity of residue 50, whereas the lowest disorder propensity in the vicinity of residue 70 is found in variants QKV06506.1 and QKQ29929.1. Finally, although variant QMU91370.1 is almost indistinguishable from variant QJS56890.1 within the first 40 residues, its intrinsic disorder predisposition in the vicinity of residue 70 is one of the lowest among all the proteins analysed in this study. Interestingly, comparison of the Fig. 9 (**A**) and Fig. 9 (**B**) show that the variability in the disorder predisposition between many variants of the protein ORF8 from SARS-CoV-2 isolates is noticeably greater than that between the reference ORF8 from SARS-CoV-2 and ORF8 proteins from Bat-CoV RaTG13 and Pangolin-CoV.

### 3.3. Possible Flow of Mutations over ORF8

Here we present five different possible mutation flows according to the date of collection of the virus sample from patients [45]. Sequence homology- and amino acid compositions-based phylogenies have been drawn for the ORF8 proteins associated in each flow.

#### Flow-I

In this flow of mutations (Fig. 13), we have described the occurrence of sequences of mutations in the US sequences based on the chronological order considering the Wuhan ORF8 sequence YP_009724396 as the reference sequence. The protein sequence QMI92505.1 possesses a mutation L4F which was of neutral type with no change in polarity. However, it showed a decreasing effect on the stability of the protein. Following this sequence, another sequence, QMT48896.1, in accordance to the time scale was identified in which is a second mutation was located at D63N, which was of neutral type and change in polarity was not observed. So, this sequence accumulated two neutral mutations which may affect the function of the protein as both mutants cause a decrease in stability of the protein. As we moved along the path at the right side, we identified the sequence QMT96239.1 with another different mutation, G8R, which was of diseased-increasing type, and the polarity changed from hydrophobic to hydrophilic. A second mutation, D35Y, mutation occurred at 2nd level in the sequence QMU92030.1 i.e this sequence was found to have two mutations G8R and D35Y. Among them, one is neutral and the other is of disease-increasing type, which in combination may alter both the structure and function of the protein.

**Figure 13:**
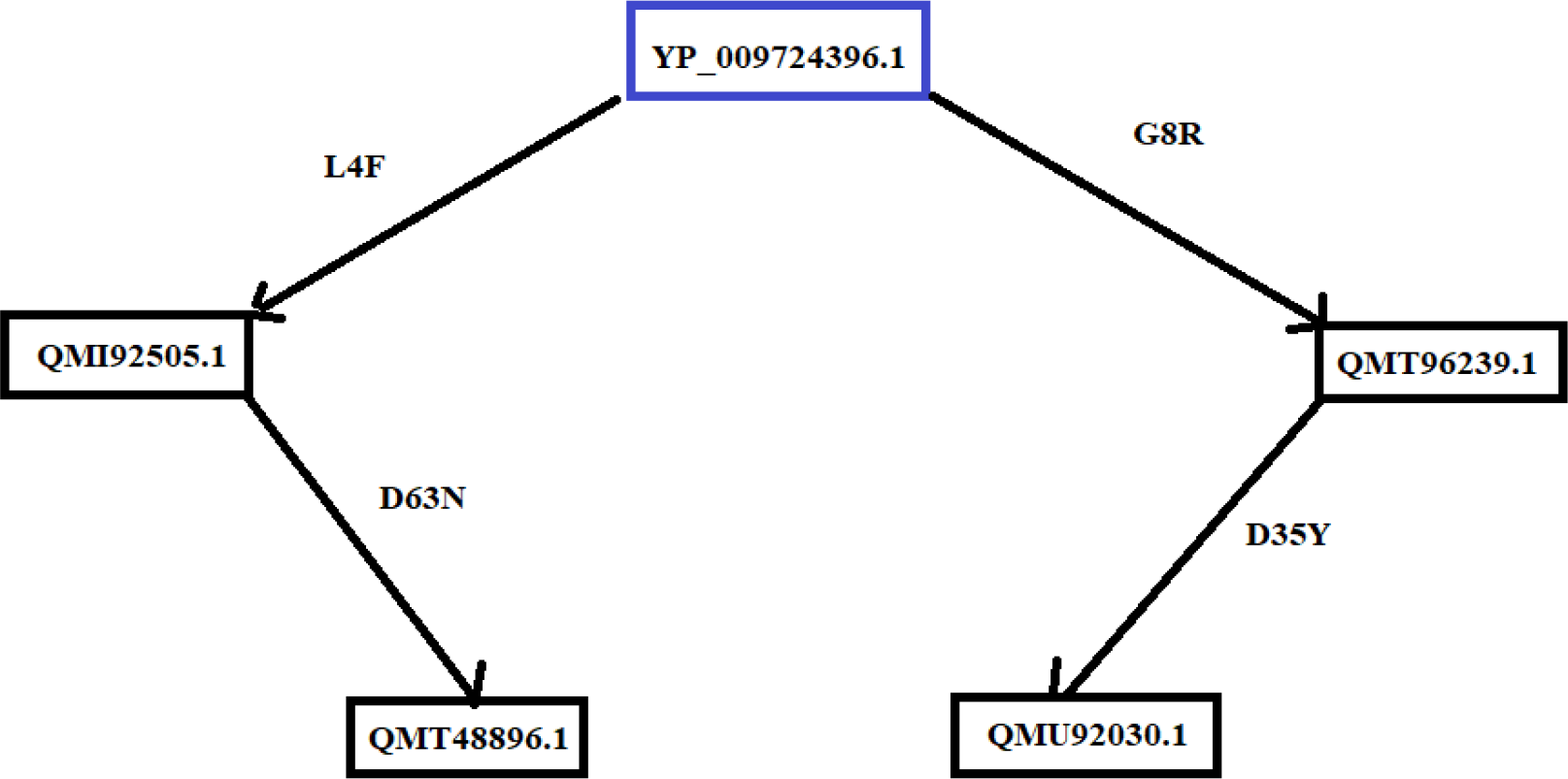
Possible flow of mutations in the ORF8 (SARS-CoV-2) sequences isolated in the US.

In order to support these mutation flows, we analysed the protein sequence similarity based on phylogeny and amino acid composition. The reference ORF8 sequence YP_009724396 is found to be much more similar to the variants QMT48896.1 and QMI92505.1, which are more similar to each other as depicted in the sequence based phylogeny (Fig. 14 (left)). This sequence based similarity of the ORF8 proteins QMT48896.1 and QMI92505.1 is illustrated in the chronology of mutations as shown in Fig. 13. Similarly, the mutation flow of the sequences QMT96239.1 and QMU92030.1 is supported by the respective sequence-based similarity.

**Figure 14:**
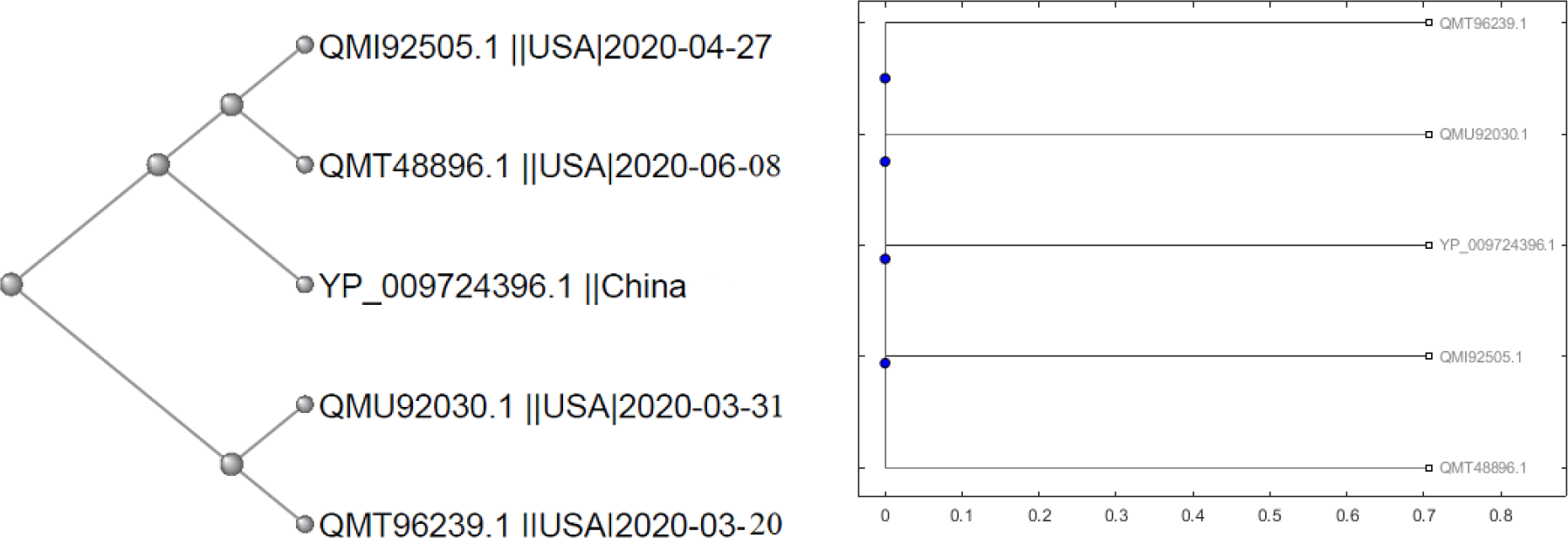
Phylogenetic relationship based on amino acid sequence similarity (left) and amino acid composition (right) of ORF8 proteins of SARS-CoV-2

The network of five ORF8 protein variants from the US is justified based on similar amino acid compositions/conservations across the five sequences as shown in Fig. 14 (right).

#### Flow-II

In this flow of mutations (Fig. 15), we observed one sequence with first-order mutations, i.e where only one mutation accumulated in the sequence. Additionally, four sequences (all are from the US) were identified with second order mutations stating that four sequences were found to have two mutations.

**Figure 15:**
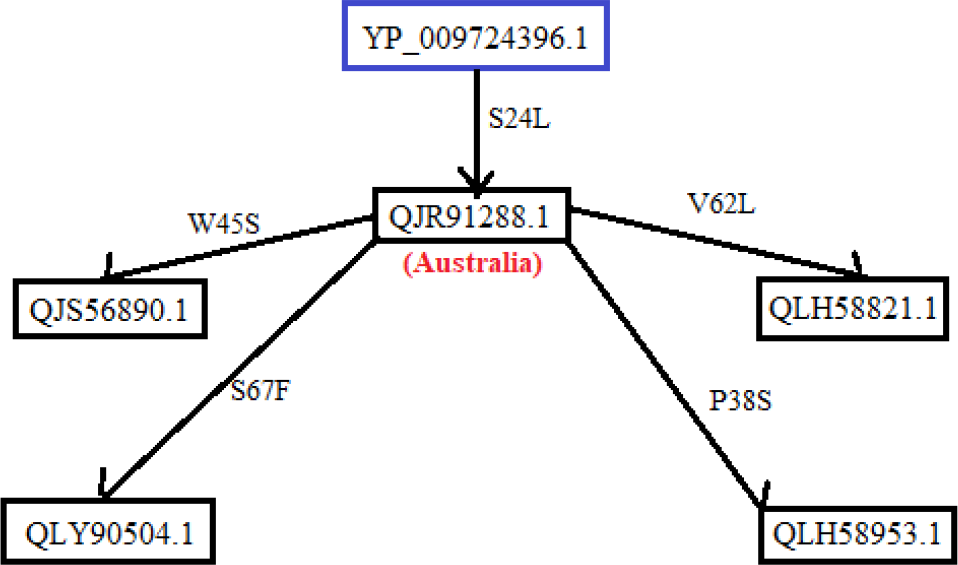
Possible flow of mutations in the ORF8 (SARS-CoV-2) sequences from the US and Australia

The protein sequence QLY90504.1 possesses a second mutation at position 67, which changed the hydrophilic serine (S) to the hydrophobic phenylalanine (F), so it may account for disrupting the ionic interactions. As it is a neutral mutation, the sequence accumulated two neutral mutations. The protein sequence QLH58953.1 acquired a second mutation, P38S, which was found to be of disease-increasing type and the polarity also changed from hydrophobic to hydrophilic, thus indicating these mutations may have some significant importance. The protein sequence QLH58821.1 possesses a second mutation, V62L, which was found to be of neutral type with no change in polarity. Here, this sequence accumulated two neutral mutations, which may account for some functional changes.

By comparing both the sequence-based phylogeny (Fig. 16 (left)) and amino acid conservation-based phylogeny (Fig. 16 (right)), we found that according to sequence-based phylogeny the Australian sequence is closely related to the ORF8 Wuhan sequence.

**Figure 16:**
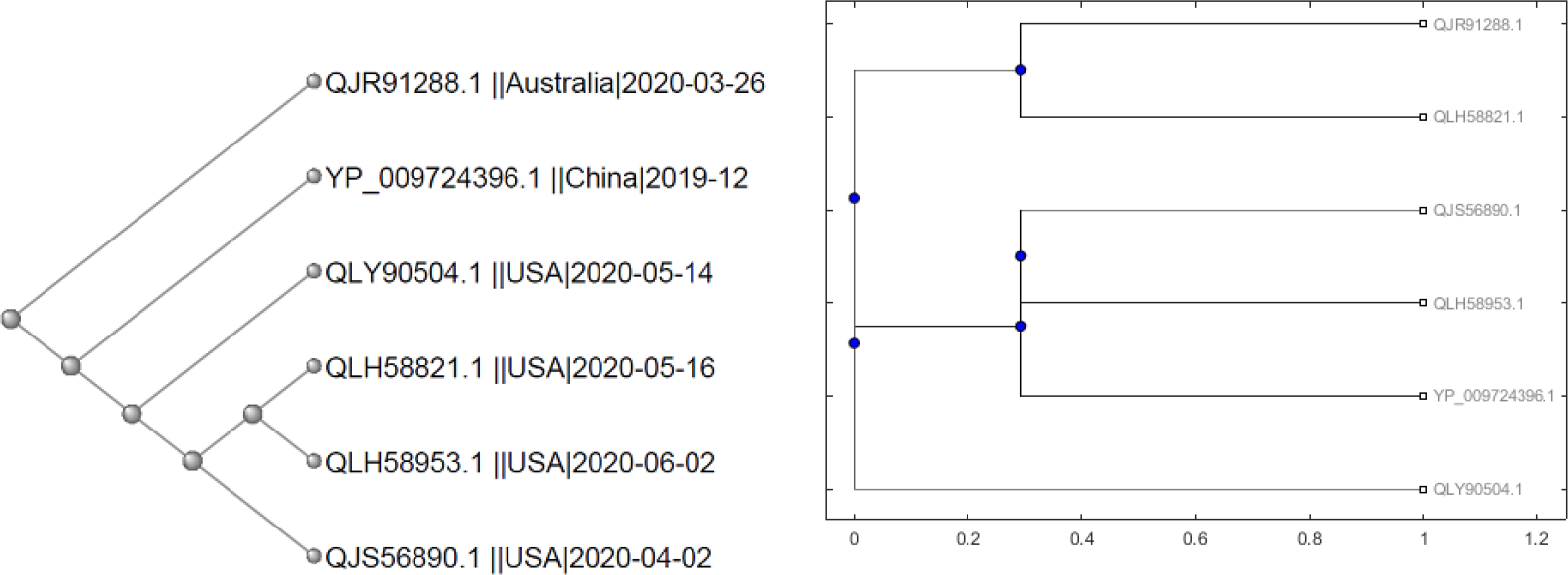
Phylogenetic relationship based on amino acid sequence similarity (left) and amino acid composition (right) of SARS-CoV-2 ORF8 proteins.

However, according to the pathway, it should be closely related to both the Wuhan sequence and the sequences having second order mutation. This can be attributed to the presence of 119 amino acid residues instead of 121 aa residues. In this case, the sequence has a two amino acid deletion, therefore, it is present at first node.

#### Flow-III

Here we analysed the US sequences considering the Wuhan sequence (YP_009724396.1) as the reference and found one sequence, QKC05159.1, with a single mutation and seven sequences with two mutations each (Fig. 17). The first sequence, QKC05159.1, contained the L84S mutation (strain determining mutation), which was of neutral type. However, the polarity changed from hydrophobic to hydrophilic, which may account for some significant change of function. The sequences that accumulated 2nd mutation along with L84S are as follows:

**Figure 17:**
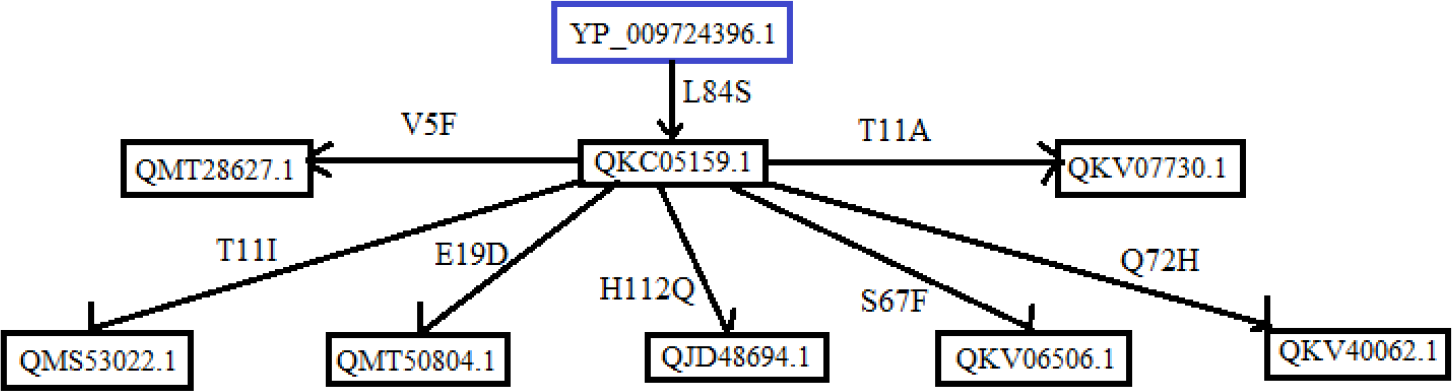
Possible flow of mutations in the ORF8 (SARS-CoV-2) sequences from the US

- **QMT28672.1**: This sequence possesses a second mutation V5F, which was predicted to be of neutral type with no change in polarity, hence this sequence acquired two neutral mutations, and together these mutations may alter the function of the protein.
- **QMS53022.1**: This protein sequence acquired a second mutation at position 11, which changed the hydrophilic amino acid threonine (T) to hydrophobic amino acid isoleucine (I) therefore affecting the ionic interactions. This mutation was found to be a diseased-increasing type, so it may affect the structure of the protein.
- **QMT50804.1**: This sequence gained a second mutation, E19D, which was predicted to be of disease-increasing type with no change in polarity. The sequence first accumulated a neutral mutation then a disease-increasing mutation, signifying that these mutations may have some functional importance.
- **QJD48694.1**: H112Q occurred as second mutation in this sequence, which was found to be of disease-increasing type with no change in polarity. Consequently, these mutations may contribute to immune evasion property of the virus.
- **QKV06506.1**: This sequence possesses the S67F mutation, which was predicted to be of neutral type, which changed the hydrophilic serine (S) to the hydrophobic phenylalanine (F), thus interfering with the ionic interactions that may increase or decrease the affinity of the viral protein for a particular host cell protein.
- **QKV40062.1**: This sequence acquired a second mutation at Q72H, which was found to be a neutral mutation and no change in polarity was observed. As this sequence accumulated two neutral mutations, it can be assumed that neutral mutations also have a significant importance.
- **QKV07730.1**: The T11A mutation occurred as the second mutation in this sequence, which was predicted to be of disease-increasing type and the polarity was changed from hydrophilic to hydrophobic, hence the structure and function of the protein are expected to differ.

From the sequence based phylogeny (Fig. 18 (left)) it was observed that the Wuhan sequence was the first to originate. Although, QKC05159.1 is the first sequence in our flow considering the time, it was found that in the phylogenetic tree it is present at fourth node instead of second node, which is probably due to the presence of ambiguous mutations in this sequence. It was also determined that QKV07730.1 is very similar to QMT50804.1 and again QMT28672.1 is more similar to both of them. All the other sequences having second order mutations are closely related to each other and follow the chronology.

**Figure 18:**
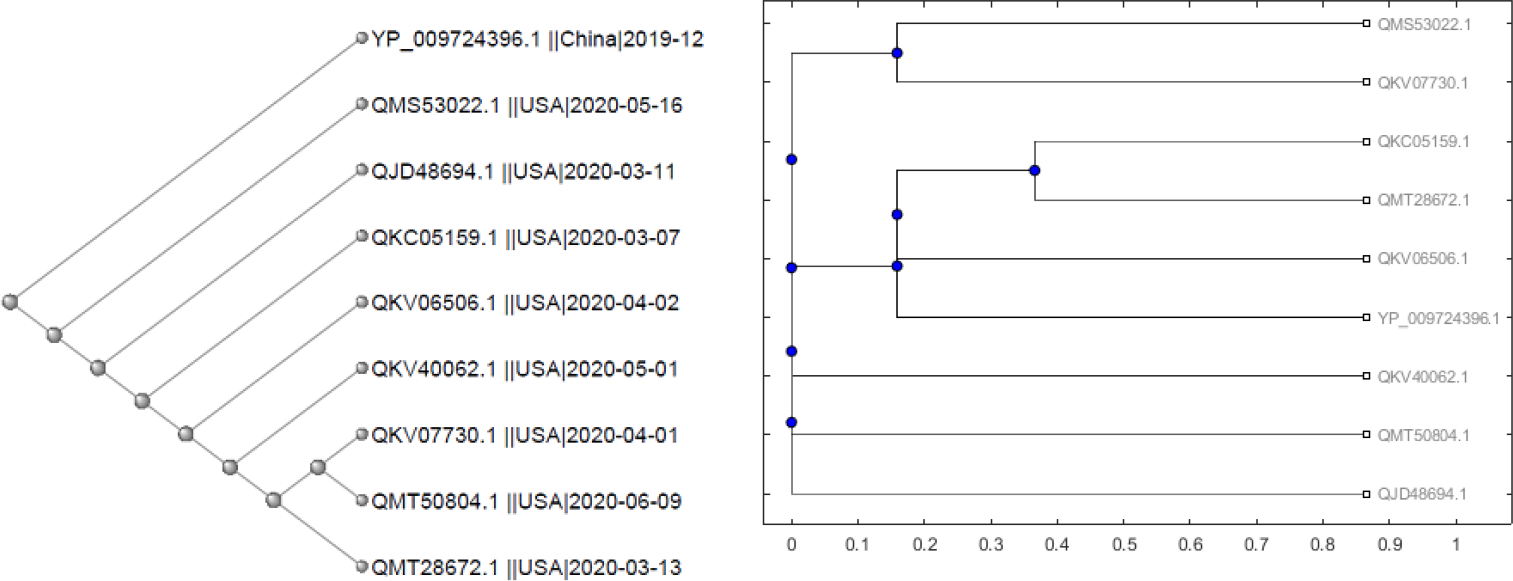
Phylogenetic relationship based on amino acid sequence similarity (left) and amino acid composition (right) ORF8 proteins of SARS-CoV-2

From the amino acid based analysis (Fig. 18(right)) it was found that the Wuhan sequence has a high conservation similarity with that of QKV06506.1, thus proving that this sequence was identified chronologically after the Wuhan sequence followed by QKC05159.1 and QMT28672.1, which again are very similar to each other.

#### Flow IV

In the possible flow of mutations (Fig. 19), we have found one sequence with a single mutation, six sequences with two mutations, and another two sequences with three mutations. The US sequence QKC05159.1 was identified to have the L84S mutation, which is a strain determining mutation and was predicted to be a neutral mutation where polarity was changed from hydrophobic to hydrophilic. The sequences that accumulated second mutations along with L84S are as following and it should be noted that the mutational accumulation occurred in a single strain:

**Figure 19:**
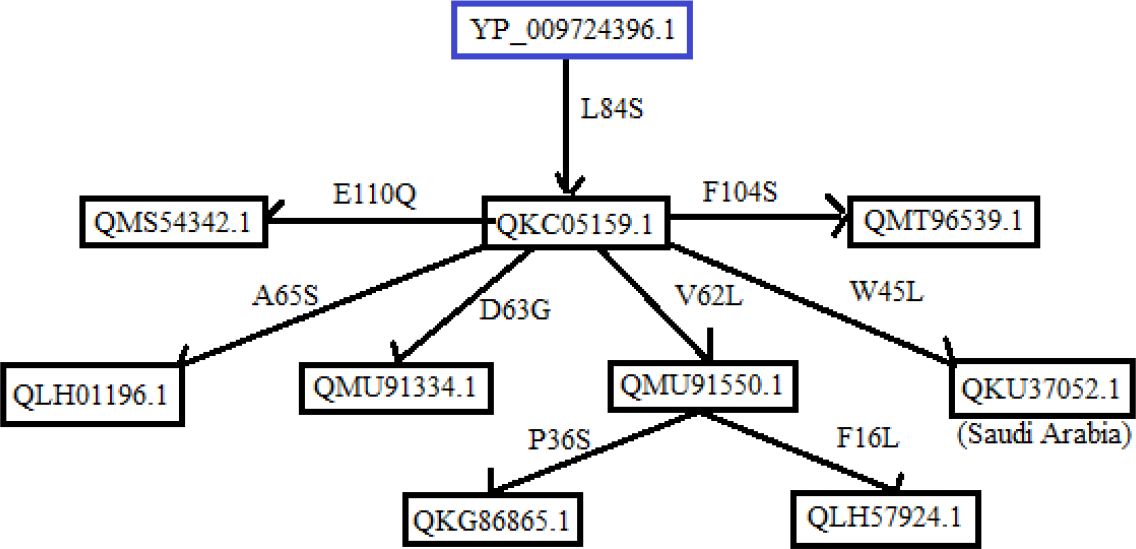
Possible flow of mutations in ORF8 (SARS-CoV-2) sequences from the US and Saudi Arabia

- **QMS54342.1**: This US sequence acquired the E110Q mutation, which was predicted to be of disease-increasing type, where no change in polarity was observed and consequently it may contribute to virulence properties of the virus.
- **QLH01196.1**: The A65S mutation occurred as a second mutation in this US sequence, which was found to be of neutral type. However, the polarity changed from hydrophobic to hydrophilic, thus potentially influencing the function of the protein.
- **QMU91334.1**: This US sequence possesses the D63G mutation, which was predicted to be of neutral type. However, the polarity changed from hydrophilic to hydrophobic so, this sequence accumulated two neutral mutations, which may allow the virus to evolve in terms of virulence.
- **QMU91550.1**: This US sequence mutated at position 62, which changed the amino acid valine (V) to lysine (L) thereby increasing the hydrophobicity of the protein. However, it was a neutral mutation even though it may influence the virulence properties of the protein. This sequence was followed by another sequence QKG86865.1 which acquired a third mutation at position 36, which changed the hydrophobic amino acid proline (P) to the hydrophilic amino acid serine (S). The mutation was neutral, thus accumulating two neutral and one disease-increasing mutations, being of significant importance for the evolution of the virus. We identified one more sequence QLH57924.1, which possesses another third mutation, F16L, which was predicted to be a neutral mutation and no polarity change was observed. This sequence acquired three neutral mutations that may promote virus survival.
- **QKU37052.1**: This sequence with the W45L mutation was reported in Saudi Arabia, which was found to be of disease-increasing type with no polarity change. Therefore, this sequence also accumulated one neutral and one disease-increasing mutation, which may affect both the structure and function of the protein.
- **QMT96539.1**: The F104S mutation was reported in the US sequence, which was found to be of disease-increasing type and the polarity changed from hydrophobic to hydrophilic. Altogether, the sequence possesses one neutral and one disease-increasing mutation that may allow the virus to acquire new properties for better survival strategies. Sequence-based phylogeny (Fig. 20 (left)) suggested that the Wuhan sequence originated first. Due to the presence of an ambiguous amino acid sequence, QKC05159.1 did not show close similarity to the Wuhan sequence.

**Figure 20:**
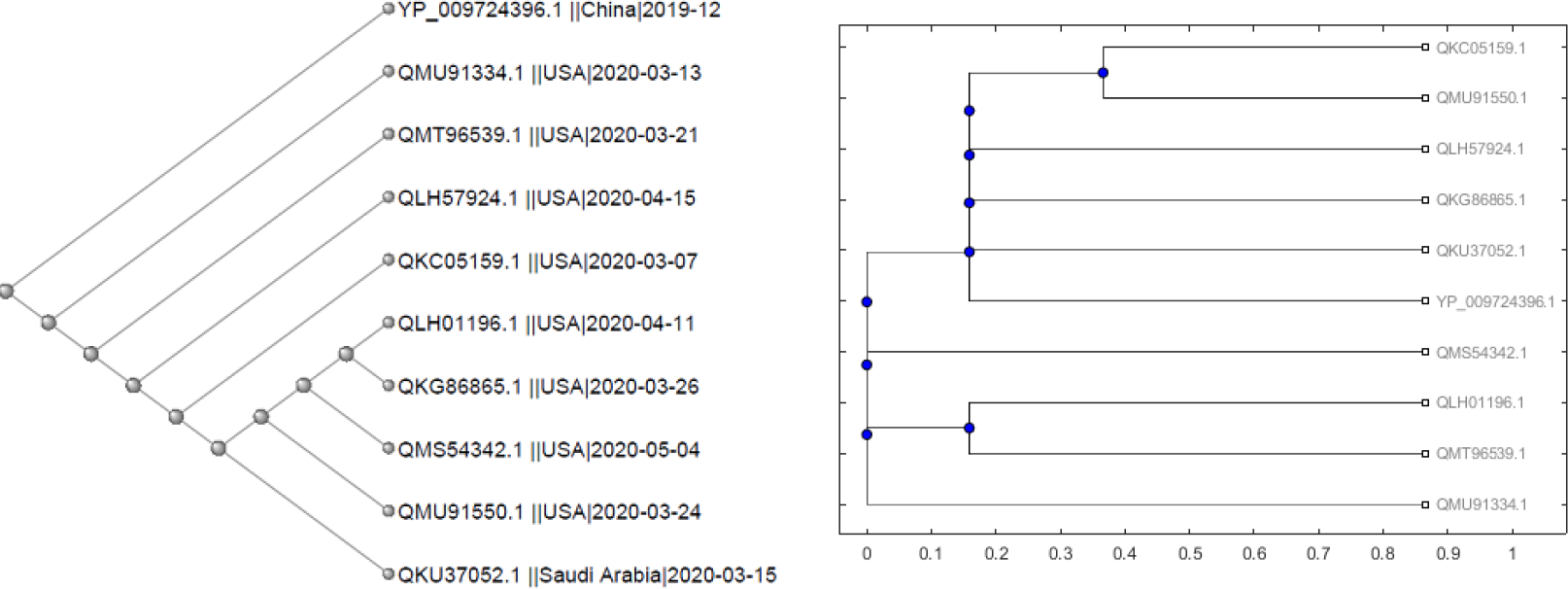
Phylogenetic relationship based on amino acid sequence similarity (left) and amino acid composition (right)of the ORF8 proteins of SARS-CoV-2

Based on sequence based phylogeny (Fig. 20 (left)) it was observed that Wuhan sequence originated first. Due to presence of ambiguous amino acids sequence QKC05159.1 was not observed in close proximity to Wuhan sequence.

QKG86865.1 and QLH57924.1 were found to have third order mutations and they are assumed to be closely related by the flow and the same has been supported by amino acid conservation-based phylogeny (Fig. 20 (right)).

#### Flow-V

QJR88780.1 (Australia) possesses the mutation L84S with reference to the Wuhan ORF8 sequence YP_009724396.1 (Fig. 21). Another sequence QJR88936.1 was reported, which possesses a second mutation, V62L. This mutation was predicted to be neutral with no change in polarity. However, the hydrophobicity increased.

**Figure 21:**
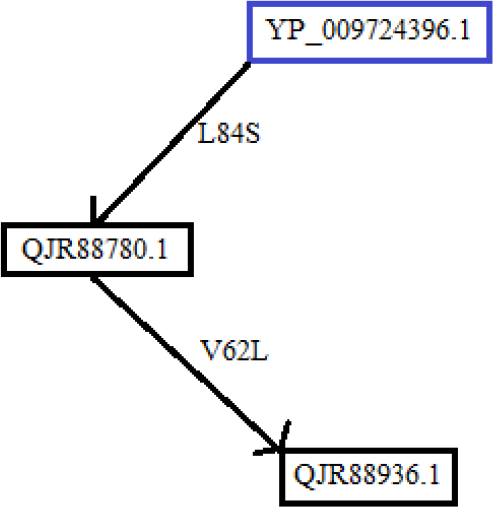
Possible flow of mutations in ORF8 (SARS-CoV-2) sequences from Australia

This sequence belongs to a particular strain and acquired two neutral mutations, indicating that these mutations may play some important role in the function of ORF8a. As can be seen from both the sequence-based phylogeny and amino acid conservation-based phylogeny (Fig. 22), the Wuhan sequence has originated earlier and the sequences QJR88780.1 and QJR88936.1 are more closely related to each other than to the Wuhan sequence as both sequences have one common mutation not present in the Wuhan sequence.

**Figure 22:**
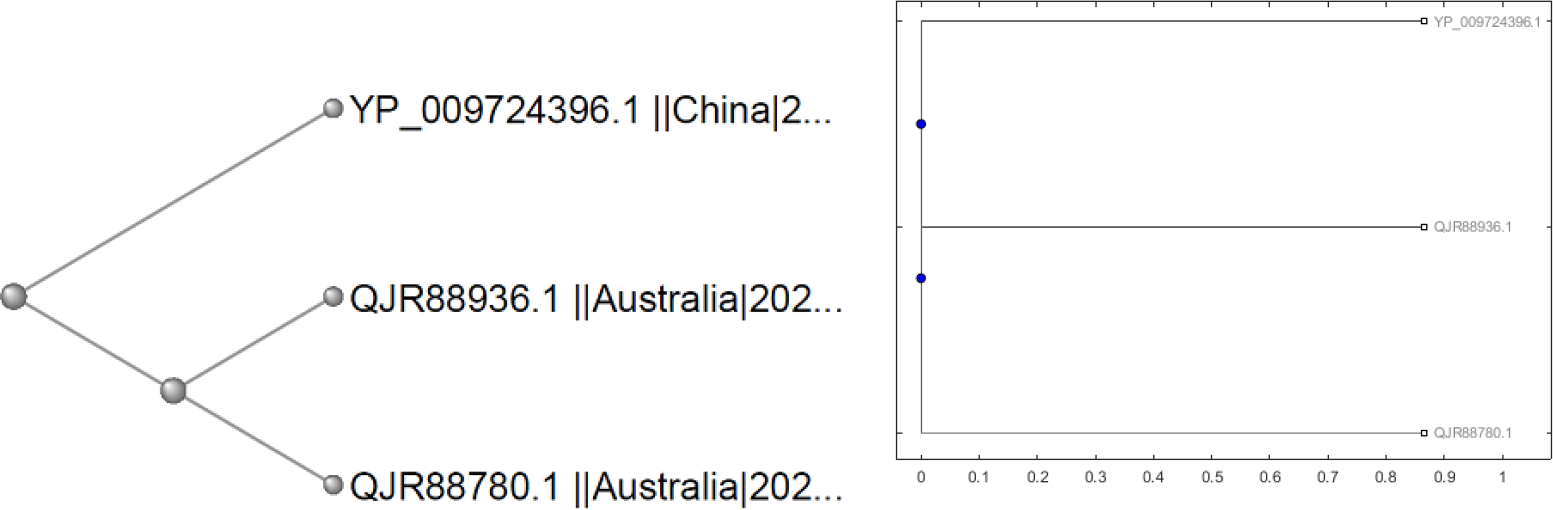
Phylogenetic relationship based on amino acid sequence similarity (left) and amino acid composition (right) of the ORF8 proteins of SARS-CoV-2

## 4. Discussion

Among SARS-CoV-2 proteins, the ORF8 accessory protein is crucial because it plays a vital role in bypassing the host immune surveillance mechanism. This protein is found to have a wide variety of mutations and among them L84S (23) and S24L (7) have highest frequency of occurrence, which bears distinct functional significance as well. It has been reported that L84S and S24L show antagonistic effects on the protein folding stability of SARS-CoV-2 [44]. L84S destabilizes protein folding, therefore up-regulating the host-immune activity and S24L favours folding stability positively, thus enhancing the functionality of ORF8 protein. L84S is already established as a strain determining mutation and since according to our studies both L84S and S24L do not occur together in a single sequence of SARS-CoV-2 ORF8 protein, it is proposed that virus with the S24L mutation is a new strain altogether. We also observed that hydrophobic to hydrophobic mutations are dominant in the D1 domain. Therefore, hydrophobicity is an important property for the N-terminal signal peptide. However, in the D2 domain, hydrophobic to hydrophilic mutations are observed more frequently, consequently making the ionic interactions more favourable and allowing the protein to evolve in terms of better efficacy in pathogenicity.

The ORF8 sequence of SARS-CoV-2 shows 93% similarity with the Bat-CoV RaTG13 and 88% similarity with that of the Pangolin-CoV ORF8. Thus, the ORF8 protein of SARS-CoV-2 can be considered as a valuable candidate for evolutionary deterministic studies and for the identification of the origin of SARS-CoV-2 as a whole. We also analysed a wide variety of mutations in the SARS-CoV-2 ORF8, where we compared it with the ORF8 of Bat-CoV RaTG13 and Pangolin-CoV in relation to charge and hydrophobicity perspectives and we found that the Bat-CoV RaTG13 ORF8 protein exhibits exactly the same properties as that of the SARS-CoV-2 ORF8 protein, whereas the properties of the Pangolin-CoV ORF8 are relatively less similar to the SARS-CoV-2 ORF8. Furthermore, to study the evolutionary nature of mutations in the ORF8, we aligned three bat sequences and found that two of them were exactly the same and there were only six amino acid differences in the third with respect to the other two sequences. So, only two variants were identified for Bat-CoV. Therefore, it shows that the rate of occurrence of mutations is slow in the Bat-CoV RaTG13 ORF8. However, for pangolins no differences were observed among four Pangolin-CoV ORF8 sequences and therefore only a single variant of ORF8 is present. Based on sequence alignment, biochemical characteristics and secondary structural analysis, the Bat-CoV, the Pangolin-CoV and the SARS-CoV-2 ORF8 displayed a high similarity index. Additionally, in the ORF8 of SARS-CoV-2, certain mutations were found to exhibit exact reversal with respect to bats and pangolins and therefore pointing towards the genomic origin of SARS-CoV-2.

However, unlike Bat-CoV and Pangolin-CoV, the mutational distribution of the ORF8 (SARS-CoV-2) is widespread ranging from the position 3 to 121, having no defined conserved region. This surprises the scientific community enormously. Further this property differentiates the SARS-CoV-2 ORF8 from that of Bat-CoV and Pangolin-CoV, thus raising the question over the natural trail of evolution of mutations in SARS-CoV-2.

We further predicted the types and effects of mutations of 95 sequences and grouped them into four domains and found that diseased type mutations with decreasing effect on stability are more prominent. Consequently, it is hypothesized that these mutations are promoting the viral survival rate. Furthermore, we tracked the possible flow of mutations in accordance to time and geographic locations and validated our proposal with respect to sequence-based and amino acid conservation-based phylogeny and therefore putting forward the order of accumulation of mutations.

This study adumbrates a comprehensive analysis of uniqueness of SARS-CoV-2, pandemic causing virus through the light of one of its accessory proteins, the ORF8 protein. In future endeavours, more critical studies on the ORF8 protein of SARS-CoV-2 is necessary for a better understanding of the importance of high frequency mutations and their role related to the host-immune system and to more correctly validate the origin of SARS-CoV-2.

## Author Contributions

SSH conceived the problem. SG,DA, SSH, VNU examined the mutations and performed analysis. SSH, PPC, SG, DA, VNU analyzed the results. SH wrote the initial draft. SH, SG, DA, PPC, VNU, AAAA, GKA, NR, KL edited the final draft. All the authors approved the final manuscript.

## Conflict of Interests

The authors do not have any conflicts of interest to declare.

## Acknowledgement

Authors thank Prof. Bidyut Roy of *Human Genetics Unit, Indian Statistical Institute, Kolkata, Indis* for his kind support for structure predictions.

## Notes

### Competing Interest Statement

The authors have declared no competing interest.

## References

[1] H. A. Rothan, S. N. Byrareddy, The epidemiology and pathogenesis of coronavirus disease (covid-19) outbreak, Journal of autoimmunity (2020) 102433.

[2] A. Zumla, M. S. Niederman, The explosive epidemic outbreak of novel coronavirus disease 2019 (covid-19) and the persistent threat of respiratory tract infectious diseases to global health security, Current Opinion in Pulmonary Medicine (2020).

[3] M. U. Kraemer, C.-H. Yang, B. Gutierrez, C.-H. Wu, B. Klein, D. M. Pigott, L. Du Plessis, N. R. Faria, R. Li, W. P. Hanage, et al., The effect of human mobility and control measures on the covid-19 epidemic in china, Science 368 (6490) (2020) 493–497.

[4] W. H. Organization, et al., Coronavirus disease 2019 (covid-19): situation report, 82 (2020).

[5] M. Ceccarelli, M. Berretta, E. V. Rullo, G. Nunnari, B. Cacopardo, Editorial–differences and similarities between severe acute respiratory syndrome (sars)-coronavirus (cov) and sars-cov-2. would a rose by another name smell as sweet?, European review for medical and pharmacological sciences 24 (2020) 2781–2783.

[6] J. Xu, S. Zhao, T. Teng, A. E. Abdalla, W. Zhu, L. Xie, Y. Wang, X. Guo, Systematic comparison of two animal-to-human transmitted human coronaviruses: Sars-cov-2 and sars-cov, Viruses 12 (2) (2020) 244.

[7] Y.-Z. Zhang, E. C. Holmes, A genomic perspective on the origin and emergence of sars-cov-2, Cell (2020).

[8] Z. Shen, Y. Xiao, L. Kang, W. Ma, L. Shi, L. Zhang, Z. Zhou, J. Yang, J. Zhong, D. Yang, et al., Genomic diversity of sars-cov-2 in coronavirus disease 2019 patients, Clinical Infectious Diseases (2020).

[9] J.-Y. Li, C.-H. Liao, Q. Wang, Y.-J. Tan, R. Luo, Y. Qiu, X.-Y. Ge, The orf6, orf8 and nucleocapsid proteins of sars-cov-2 inhibit type i interferon signaling pathway, Virus research (2020) 198074.

[10] D. Kim, J.-Y. Lee, J.-S. Yang, J. W. Kim, V. N. Kim, H. Chang, The architecture of sars-cov-2 transcriptome, Cell (2020).

[11] Y. Zhang, J. Zhang, Y. Chen, B. Luo, Y. Yuan, F. Huang, T. Yang, F. Yu, J. Liu, B. Liu, et al., The orf8 protein of sars-cov-2 mediates immune evasion through potently downregulating mhc-i, bioRxiv (2020).

[12] S. Mohammad, A. Bouchama, B. M. Alharbi, M. Rashid, T. S. Khatlani, N. S. Gaber, S. S. Malik, Sars-cov-2 orf8 and sars-cov orf8ab: Genomic divergence and functional convergence (2020).

[13] R. A. Khailany, M. Safdar, M. Ozaslan, Genomic characterization of a novel sars-cov-2, Gene reports (2020) 100682.

[14] Y. Tan, T. Schneider, M. Leong, L. Aravind, D. Zhang, Novel immunoglobulin domain proteins provide insights into evolution and pathogenesis of sars-cov-2-related viruses, Mbio 11 (3) (2020).

[15] S. Chen, X. Zheng, J. Zhu, R. Ding, Y. Jin, W. Zhang, H. Yang, Y. Zheng, X. Li, G. Duan, Extended orf8 gene region is valuable in the epidemiological investigation of sars-similar coronavirus, The Journal of Infectious Diseases (2020).

[16] I. Alam, A. K. Kamau, M. Kulmanov, S. T. Arold, A. T. Pain, T. Gojobori, C. M. Duarte, Functional pangenome analysis provides insights into the origin, function and pathways to therapy of sars-cov-2 coronavirus (2020).

[17] I. Alam, A. A. Kamau, M. Kulmanov, L. Jaremko, S. T. Arold, A. Pain, T. Gojobori, C. M. Duarte, Functional pangenome analysis shows key features of e protein are preserved in sars and sars-cov-2, Frontiers in Cellular and Infection Microbiology 10 (2020) 405.

[18] J. Wu, X. Yuan, B. Wang, R. Gu, W. Li, X. Xiang, L. Tang, H. Sun, Severe acute respiratory syndrome coronavirus 2: From gene structure to pathogenic mechanisms and potential therapy, Frontiers in microbiology 11 (2020).

[19] S. R. Schaecher, A. Pekosz, Sars coronavirus accessory gene expression and function, in: Molecular Biology of the SARS-Coronavirus, Springer, 2010, pp. 153–166.

[20] A. Parlikar, K. Kalia, S. Sinha, S. Patnaik, N. Sharma, S. G. Vemuri, G. Sharma, Understanding genomic diversity, pan-genome, and evolution of sars-cov-2, PeerJ 8 (2020) e9576.

[21] M. D. Park, Immune evasion via sars-cov-2 orf8 protein? (2020).

[22] S. Kumar, R. Nyodu, V. K. Maurya, S. K. Saxena, Host immune response and immunobiology of human sars-cov-2 infection, in: Coronavirus Disease 2019 (COVID-19), Springer, 2020, pp. 43–53.

[23] S.-C. Sung, C.-Y. Chao, K.-S. Jeng, J.-Y. Yang, M. M. Lai, The 8ab protein of sars-cov is a luminal er membrane-associated protein and induces the activation of atf6, Virology 387 (2) (2009) 402–413.

[24] M. Frieman, R. Baric, Mechanisms of severe acute respiratory syndrome pathogenesis and innate immunomodulation, Microbiology and Molecular Biology Reviews 72 (4) (2008) 672–685.

[25] The Mathworks, Inc., Natick, Massachusetts, MATLAB version 9.3.0.713579 (R2020a) (2020).

[26] X. Wang, Q. Zhou, Y. He, L. Liu, X. Ma, X. Wei, N. Jiang, L. Liang, Y. Zheng, L. Ma, et al., Nosocomial outbreak of 2019 novel coronavirus pneumonia in wuhan, china, European Respiratory Journal (2020).

[27] R. Cotton, Current methods of mutation detection, Mutation Research/Fundamental and Molecular Mechanisms of Mutagenesis 285 (1) (1993) 125–144.

[28] G. M. Boratyn, J. Thierry-Mieg, D. Thierry-Mieg, B. Busby, T. L. Madden, Magic-blast, an accurate rna-seq aligner for long and short reads, BMC bioinformatics 20 (1) (2019) 1–19.

[29] F. Madeira, Y. M. Park, J. Lee, N. Buso, T. Gur, N. Madhusoodanan, P. Basutkar, A. R. Tivey, S. C. Potter, R. D. Finn, et al., The embl-ebi search and sequence analysis tools apis in 2019, Nucleic acids research 47 (W1) (2019) W636–W641.

[30] E. Capriotti, R. B. Altman, Y. Bromberg, Collective judgment predicts disease-associated single nucleotide variants, BMC genomics 14 (S3) (2013) S2.

[31] E. Capriotti, P. Fariselli, R. Casadio, I-mutant2. 0: predicting stability changes upon mutation from the protein sequence or structure, Nucleic acids research 33 (suppl_2) (2005) W306–W310.

[32] D. Xu, Y. Zhang, Ab initio protein structure assembly using continuous structure fragments and optimized knowledge-based force field, Proteins: Structure, Function, and Bioinformatics 80 (7) (2012) 1715–1735.

[33] D. Xu, Y. Zhang, Toward optimal fragment generations for ab initio protein structure assembly, Proteins: Structure, Function, and Bioinformatics 81 (2) (2013) 229–239.

[34] D. Peacock, D. Boulter, Use of amino acid sequence data in phylogeny and evaluation of methods using computer simulation, Journal of molecular biology 95 (4) (1975) 513–527.

[35] J. A. Irving, R. N. Pike, A. M. Lesk, J. C. Whisstock, Phylogeny of the serpin superfamily: implications of patterns of amino acid conservation for structure and function, Genome research 10 (12) (2000) 1845–1864.

[36] F. Austerlitz, O. David, B. Schaeffer, K. Bleakley, M. Olteanu, R. Leblois, M. Veuille, C. Laredo, Dna barcode analysis: a comparison of phylogenetic and statistical classification methods, BMC bioinformatics 10 (S14) (2009) S10.

[37] S. S. Hassan, P. P. Choudhury, P. Basu, S. S. Jana, Molecular conservation and differential mutation on orf3a gene in indian sars-cov2 genomes, Genomics 112 (5) (2020) 3226–3237.

[38] S. S. Hassan, P. P. Choudhury, B. Roy, Molecular conservation and differential mutation on orf3a gene in indian sars-cov2 genomes, Genomics 112 (6) (2020) 3890–3892.

[39] S. S. Hassan, P. P. Choudhury, B. Roy, S. S. Jana, Molecular conservation and differential mutation on orf3a gene in indian sars-cov2 genomes, Genomics 112 (6) (2020) 4622–4627.

[40] Z. Obradovic, K. Peng, S. Vucetic, P. Radivojac, A. K. Dunker, Exploiting heterogeneous sequence properties improves prediction of protein disorder, Proteins: Structure, Function, and Bioinformatics 61 (S7) (2005) 176–182.

[41] F. Meng, V. N. Uversky, L. Kurgan, Comprehensive review of methods for prediction of intrinsic disorder and its molecular functions, Cellular and Molecular Life Sciences 74 (17) (2017) 3069–3090.

[42] Z.-L. Peng, L. Kurgan, Comprehensive comparative assessment of in-silico predictors of disordered regions, Current Protein and Peptide Science 13 (1) (2012) 6–18.

[43] X. Fan, L. Kurgan, Accurate prediction of disorder in protein chains with a comprehensive and empirically designed consensus, Journal of Biomolecular Structure and Dynamics 32 (3) (2014) 448–464.

[44] A. H. Rad, A. D. McLellan, Implications of sars-cov-2 mutations for genomic rna structure and host microrna targeting, bioRxiv (2020).

[45] S. S. Hassan, D. Attrish, S. Ghosh, P. P. Choudhury, B. Roy, Pathogenetic perspective of missense mutations of orf3a protein of sars-cov2, bioRxiv (2020).

